# Inferring protein-protein interaction networks from inter-protein sequence co-evolution

**DOI:** 10.1101/034744

**Authors:** Christoph Feinauer, Hendrik Szurmant, Martin Weigt, Andrea Pagnani

## Abstract

Interaction between proteins is a fundamental mechanism that underlies virtually all biological processes. Many important interactions are conserved across a large variety of species. The need to maintain interaction leads to a high degree of co-evolution between residues in the interface between partner proteins. The inference of protein-protein interaction networks from the rapidly growing sequence databases is one of the most formidable tasks in systems biology today. We propose here a novel approach based on the *Direct-Coupling Analysis* of the co-evolution between inter-protein residue pairs. We use ribosomal and trp operon proteins as test cases: For the small resp. large ribosomal subunit our approach predicts protein-interaction partners at a true-positive rate of 70% resp. 90% within the first 10 predictions, with areas of 0.69 resp. 0.81 under the ROC curves for all predictions. In the trp operon, it assigns the two largest interaction scores to the only two interactions experimentally known. On the level of residue interactions we show that for both the small and the large ribosomal subunit our approach predicts interacting residues in the system with a true positive rate of 60% and 85% in the first 20 predictions. We use artificial data to show that the performance of our approach depends crucially on the size of the joint multiple sequence alignments and analyze how many sequences would be necessary for a perfect prediction if the sequences were sampled from the same model that we use for prediction. Given the performance of our approach on the test data we speculate that it can be used to detect new interactions, especially in the light of the rapid growth of available sequence data.

## Introduction

Proteins are the major work horses of the cell. Being part of all essential biological processes, they have catalytic, structural, transport, regulatory and many other functions. Few proteins exert their function in isolation. Rather, most proteins take part in concerted physical interactions with other proteins, forming networks of protein-protein interactions (PPI). Unveiling the PPI organization is one of the most formidable tasks in systems biology today. High-throughput experimental technologies, applied for example in large-scale yeast two-hybrid [22] analysis and in protein affinity mass-spectrometry studies [20], allowed a first partial glance at the complexity of organism-wide PPI networks. However, the reliability of these methods remains problematic due to their high false-positive and false-negative rates [5].

Given the fast growth of biological sequence databases, it is tempting to design computational techniques for identifying protein-protein interactions [19]. Prominent techniques to date include: the genomic co-localization of genes [10,18] (with bacterial operons as a prominent example), the Rosetta-stone method [28] (which assumes that proteins fused in one species may interact also in others), phylogenetic profiling [34] (which searches for the correlated presence and absence of homologs across species), and similarities between phylogenetic trees of orthologous proteins [24,33,37,43]. Despite the success of all these methods, their sensitivity is limited due to the analysis of coarse global proxies for protein-protein interaction. An approach that exploits more efficiently the large amount of information stored in multiple sequence alignments (MSA) seems therefore promising.

Recently, a breakthrough has been achieved using genomic sequences for the related problem of inferring residue contacts from sequence data alone. [11]. The so-called Direct-Coupling Analysis (DCA) [31,38] allows to construct statistical models that are able to describe the sequence variability of large MSA of homologous proteins [15]. More precisely, these models reproduce the empirically measured covariations of amino acids at residue pairs. The parameters of the models unveil networks of direct residue co-evolution, which in turn accurately predict residue-residue contacts.

These models are computationally hard to infer and several approximations have therefore been developed [2, 12, 31, 38]. While models based on the mean-field approximation borrowed from statistical physics [2, 31] are fast, approximations based on pseudo-likelihood maximization [1, 12] are more accurate and used throughout this paper.

When applied to two interacting protein families, DCA and related methods are able to detect inter-protein contacts [21, 32, 38] and thereby to guide protein complex assembly [9, 36]. This is notable since contact networks in protein complexes are strongly modular: There are many more intra-protein contacts than inter-protein contacts. Moreover, DCA helps to shed light on the sequence-based mechanisms of PPI specificity [6, 7, 35].

Here we address an important question: Is the strength of inter-protein residue-residue co-evolution sufficient to *discriminate interacting from non-interacting pairs of protein families*, i.e. to infer PPI networks from sequence information? A positive answer would lever the applicability of these statistical methods from structural biology (residue contact map inference) to systems biology (PPI network inference). An obvious problem in this context is the sparsity of PPI networks, illustrated by the bacterial ribosomal subunits used in the following, cf. Fig. 1: The small subunit contains 20 proteins and 21 protein-protein interfaces (11% of all 190 possible pairs). In the large subunit, 29 proteins form 29 interfaces (7% of all 406 pairs). We see that while the number of potential PPI between *N* proteins is 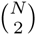, the number of real PPI grows only linearly as 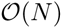. Furthermore, the number of potentially co-evolving residue-residue contacts across interfaces is much smaller than the number of intra-protein contacts. In the case of ribosomes, only 5.8% of all contacts in the small subunit are inter-protein contacts. In the large subunit this fraction drops down to 4.5%. So the larger the number of proteins, the more our problem resembles the famous search of a needle in a haystack. The noise present in the large number of non-interacting protein family pairs might exceed the co-evolutionary signal of interacting pairs.

**Figure 1:**
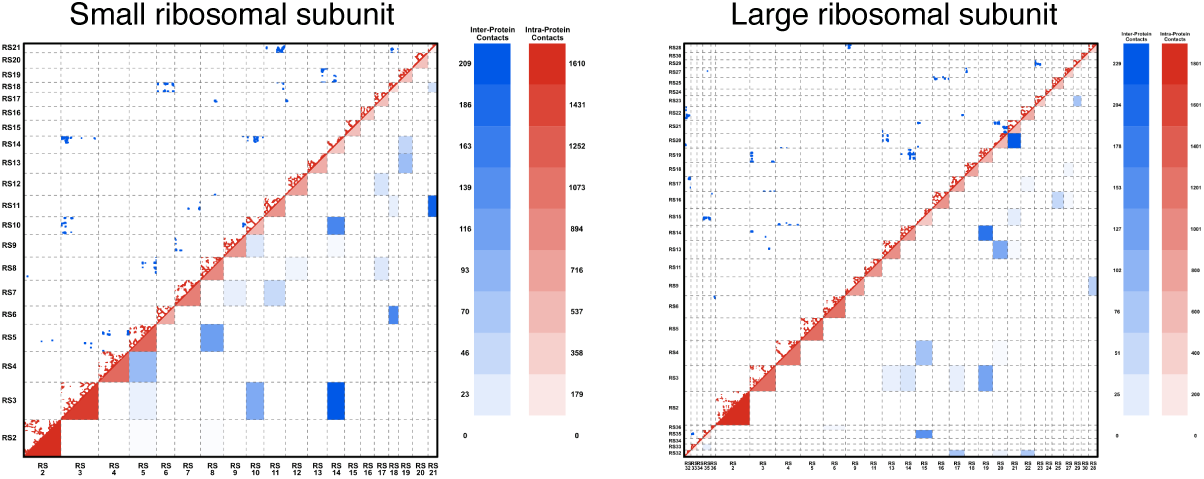
Contact map and protein-protein interaction network of small and large ribosomal subunits. The contact map and the protein-protein interaction network for **A** the small ribosomal subunit and **B** the large ribosomal subunit (proteins only), using a distance cutoff of 8Å between heavy atoms. The upper diagonal part shows the contact map, with red dots indicating intra-protein contacts, and blue dots inter-protein contacts. The lower triangular part shows the coarse graining into the corresponding protein-protein interaction networks, with the color levels indicating the number of intra-resp. inter-protein contacts, cf. the scales. The sparse character of both the contact network and the interaction network is clearly visible.

It should also be mentioned that the ribosomal structure relies on the existence of ribosomal RNA, which is not included in our analysis. We therefore expect many of the small PPI interfaces to be of little importance for the ribosomal stability and that only large interfaces constrain sequence evolution and thus become detectable by co-evolutionary studies.

Ribosomal proteins and their interactions are essential and thus conserved across all bacteria, and it appears reasonable to wonder whether this makes them a specialized example of a protein complex more amenable to co-evolutionary bias. As a second and smaller interaction network, we therefore considered the enzymes of the tryptophan biosynthesis pathway comprising a set of seven proteins in which only two pairs are known to interact (PDB-ID 1qdl for the TrpE-TrpG complex [27] and 1k7f for the TrpA-TrpB complex [40]). Also here the PPI network is very sparse; most pairs are not known to interact, but might show some degree of coordinated evolution due to the fact that in many organisms these genes show a common spatial co-localization in a single operon and also due to a number of gene fusion events, cf. the discussion below. While widespread, the tryptophan biosynthesis pathway is not essential for viability when environmental tryptophan is present.

In this paper we report the excellent performance of DCA in the prediction of protein-protein interaction partners in the systems tested. In a first step, we analyze the performance on data from an artificial model. This allows for a systematic analysis of the performance of different approaches and of the influence of the number of sequences in the alignment. With this artificial data set we are able to establish a lower-bound on the number of sequences that would make our predictions on the PPI scale completely accurate if the generating model was the same model we use for inference. Given the growth-rate of current protein sequence databases (notably UniProt [8]), we expect that such a lower bound could be met in few years. In a second step, we apply the method to the proteins of the bacterial ribosome and to the proteins of the trp operon, and show that the results obtained for simulated data translate well to the biological sequences of this test-set.

## Materials and Methods

The goal of the present work is to analyze each of the 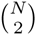 possible pairs of multiple sequence alignments from a given set of *N* single-protein family alignments, and to extract a pairwise score that measures the co-evolution between the proteins in the alignments. A high co-evolutionary score is then taken as a proxy for interaction. In the spirit of [14] we describe in this section consecutively the *data generation and matching*, the *model* used for analyzing data and the *inference and scoring* mechanism.

### Data extraction and matching for the ribosomal and trp operon proteins

The input data is given by *N* multiple sequence alignments *D_p_* consisting of *M_p_* sequences of length *L_p_* for every protein family *p*. These alignments are extracted from UniProt [8] using standard bioinformatics tools, in particular Mafft [25] and HMMer [16] (*cf*. Section 1 in S1 Text for details on the extraction pipeline and Tables S1,S2 in S1 Text listing the values of *N*, *M_p_, L_p_* for ribosomal and trp-operon proteins). For the analysis, it is necessary to concatenate the MSAs of two putative co-evolving protein families. This means to create, for each pair of protein families (*p, p′*), a new alignment *D_p,p′_* of sequence length *L_p_* + *L_p_*_′_. Each line contains the concatenation of two potentially interacting proteins. More precisely, in the case where families *p* and *p′* actually interact, each line should contain a pair of interacting proteins. The general problem of producing a concatenated alignment out of single MSAs of two protein families is straightforward in two cases only: (i) we have prior knowledge which pairs of sequences represent interaction partners; (ii) no paralogs are present in the considered species (*i.e*. all species have at most a single homolog of each of the sequences to be matched). Often, as displayed schematically in Fig. 2, MSAs contain multiple protein sequences within a given species and no prior knowledge can be used to know who is (potentially) interacting with whom. In prokaryotes, interacting proteins are frequently found to be coded in joint operons. This suggests to use genomic co-localization as a matching criterion. To do so, as explained in Section 2 in S1 Text, we approximated the *genomic distance* between sequences using UniProt accession numbers. A better distance between sequences could be defined in terms of their genomic location. Unfortunately, genomic locations are available only in the context of whole genome sequencing projects. The majority of sequences in Uniprot originate from fragments or from incomplete genome sequencing projects. These difficulties lead us to content ourselves with the proxy of accession numbers.

**Figure 2:**
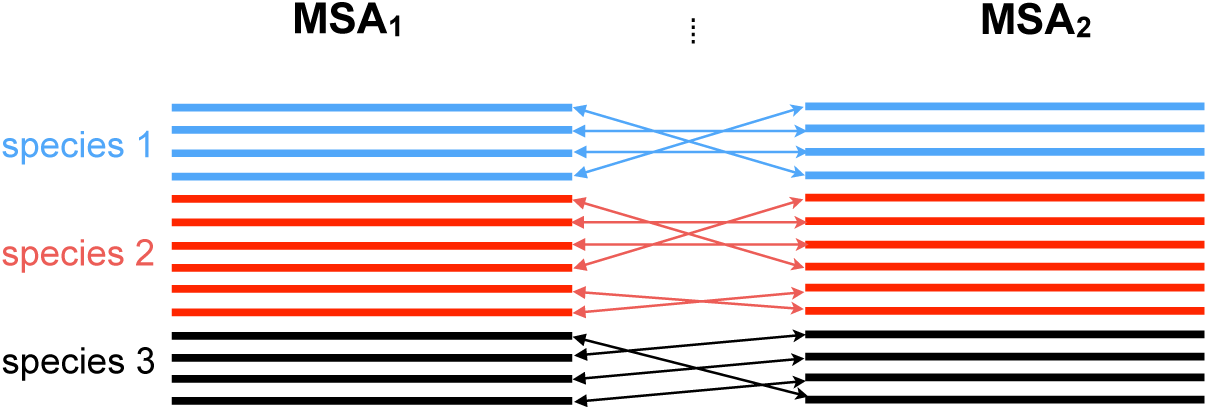
Concatenating two multiple sequence alignments. Figure Caption Sketch of the matching procedure that allows us to concatenate two different MSAs, here MSA1,MSA**2**. π represents the optimal permutation of the sequences on the second MSA computed using a standard linear programming routine.

Having defined distances between each protein pair in the MSA, we calculate the matching which minimizes the average distance between matched sequences by linear programming. Additionally, we introduce a distance threshold used to discard matched distal protein sequence pairs. The numeric value for this threshold was determined using the small ribosomal subunit as a test case.

The average number of paralogs per species varies from system to system: For both ribosomal subunits the proteins have between 1.5 and 3 paralogous sequences per genome. The trp proteins on the other hand have considerably more paralogous sequences and the number of such sequences per genome varies between 4 and 24. This means that especially in the trp operon the matching procedure has the potential to generate much larger alignments than the competing approach of excluding species with paralogous sequences. In fact, using this last approach (which corresponds to setting our threshold parameter to 0) reduces the number of sequences in the alignments on the average by about 10% for the ribosomal proteins and by about 85% for the proteins of the trp operon (see Tables S3-S7 in S1 Text).

Note that using paralogs may be dangerous since after duplication different paralogs often evolve different functions, and thus loose part of their interactions or gain others. However, our matching strategy based on genomic vicinity excludes proteins coming from isolated genes; it identifies mostly protein pairs coded in gene pairs colocalized inside an operon. It is therefore more likely that the two maintained interaction, when also the ancestral protein pair before duplication was interacting. We will show evidence that, in the interacting protein systems investigated here, this strategy leads to a reinforced coevolutionary signal. However, an independent and direct test whether protein pairs included in the alignment actually interact would constitute a big step forward.

Let us recall that the problem of finding a good matching between sequences has already been studied in the past using different strategies [6, 35]. Unfortunately, both methods are computationally too demanding to be used in a case, where hundreds or thousands of protein family pairs have to be matched.

### Statistical sequence model

Within DCA, the probability distribution over amino acid sequences *x* = (*x*_1_,…, *x_L_*) of (aligned) length *L* is modeled by a so-called Potts model, or pairwise Markov Random Field,

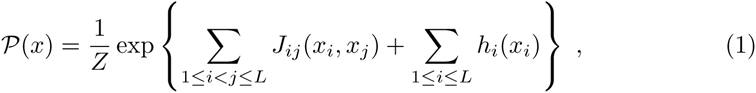

which includes statistical couplings *J_ij_*(*x_i_, x_j_*) between residue pairs and position-specific biases *h_i_*(*x_i_*) of amino-acid usage [38]. The number *Z* is the normalization constant of 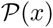, which is a probability distribution over all amino-acid sequences of length *L*. The variable *x_i_* represents the amino acid found at position *i* in the sequence and can take as values any of the *q* = 21 different possible letters in an MSA (gaps are treated as a 21st amino acid). The model parameters are inferred using MSAs of homologous proteins.

In the case of two concatenated protein sequence 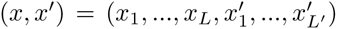, the joint probability takes the form

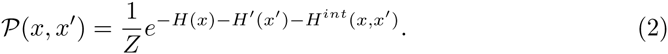

The functions *H*(*x*) and *H*′(*x*′) are the terms in the exponential in Eq. (1) referring to each single protein. The function

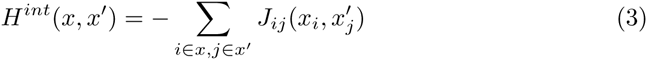

describes the co-evolutionary coupling between the two protein families. In the last expression, *x_i_* is the *i*th amino acid in sequence *x*, and 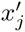 the *j*th amino acid in sequence *x′*. The sum runs over all inter-protein pairs of residue positions. The *q × q* matrices *J_ij_* in this term quantify how strongly sites between the two proteins co-evolve in order to maintain their physicochemical compatibility. The matrix contains a real number for each possible amino acid combination at sites *i* and *j* and contributes to the probability in Equation 2 depending on whether an amino acid combination is favorable or not. The strongest inter-protein couplings are enriched for inter-protein contacts [32, 38]. The same kind of model can be used to predict the interaction between more than two proteins, with a corresponding number of interaction terms. However, the number of parameters in the model is proportional to (*L*_1_ + *L*_2_ + .. + *L_N_*)^2^ for *N* proteins while the number of samples in the concatenated MSA 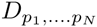 becomes smaller because one has to find matching sequences for N proteins *simultaneously*. This leads us to consider the case *N* > 2 only for artificial proteins where the total length and sample size are controllable.

### Inference and Scoring

Following [12], the parameters of the model were inferred by maximizing *pseudo-likelihood functions*. This is an alternative to directly maximizing the likelihood and considerably faster (see Section 3 in S1 Text Text for details). Given that the model is mathematically equivalent to the one used in [12] we can use the output of the algorithm (plmDCA) with default parameters as presented there directly for our purposes. This output consists of scores *F_ij_* (the average-product corrected Frobenius norm of the matrices *J_ij_*) that quantify the amount of co-evolution between sites *i* and *j* in the alignments. In order to quantify co-evolution between *proteins*, we took the *F_ij_* corresponding to inter-protein site pairs (i.e. *i* in *x* and *j* in *x′*) and calculated the mean of the 4 largest. These quantities, a real number for every protein pair, are used to rank protein-protein interaction partners. The number 4 was chosen because it performed well in the small ribosomal subunit, which we used as a test case when designing the algorithm. Subsequent tests on larger systems showed that any number between 1 and 6 performs almost equally well (see Section *A Global View* in *Results*).

### Simulated data

As the basis for the simulated data we used a fictitious protein complex consisting of 5 proteins. Each protein has a length of 53 residues. The individual contact map of each one is given by the bovine pancreatic trypsin inhibitor (PDB ID 5pti [42]), which is a small protein performing well for the prediction of internal contacts by DCA. Each *P_i_* has 551 internal contacts. Moreover, each protein interacts with two others in a circular way. The inter-protein contact matrices between *P_i_* and *P_i+_*_1_ (as well as between *P_1_* and *P_5_*) are random binary matrices with a density of 10% of the internal contacts. This models the sparsity of the inter-protein contacts as compared to the intra-protein contacts. A contact map for the artificial complex can be found in Figure S5 in S1 Text, There are no contacts between other pairs of proteins.

In order to define as realistically as possible the coupling parameters of the Potts model used for generating the artificial sequences, we used the Pfam protein family PF00014 of the pancreatic trypsin inhibitor [15]. Note that a member of this family was also used to define the structure. The couplings describing the co-evolution *within* the single proteins were directly extracted from the Pfam MSA using DCA. For the couplings corresponding to the co-evolution *between* the proteins, we used a random subset of the internal parameters and used them to couple sites that are in contact according the contact map as defined above. Non-contacting pairs of sites remain uncoupled between artificial proteins. Using this model, a joint MSA *D*_12345_ of sequences of length 265 = 5 × 53 was generated using standard MC simulations.

The process of defining the contact map, choosing the parameters and generating the sequences is described in Section 5 S1 Text.

## Results and Discussion

### Testing the approach using simulated data

As a first test of our approach, we use *simulated data* generated by Monte Carlo (MC) sampling of a Potts model of the form of Eq. (2), cf. *Materials and Methods*.

The main simplifying assumptions in this context are: (i) We assume intra- and inter-protein co-evolution strengths to be the same. (ii) We assume the distribution of inter-protein residues contacts within the possible contacts to be random. (iii) We assume the sequences to be identically and independently distributed according to our model. This model includes the assumption that non-contacting sites have zero couplings. The number of artificial sequences needed for a good performance of our method should therefore be taken at most as a lower bound for the number of biological sequences needed for a comparable performance.

In panel A of Fig. 3 we show the architecture of our artificial protein complex. It is composed of five fictitious, structurally identical proteins *P*_1_,…, *P*_5_, each one consisting of 53 residues. In order to simulate co-evolution between the proteins, we generate a *joint* MSA *D*_12345_ for all 5 proteins with a model that contains couplings between inter-protein site pairs. These couplings are modeled in a way to resemble couplings inferred from real proteins (see *Materials and Methods*).

**Figure 3:**
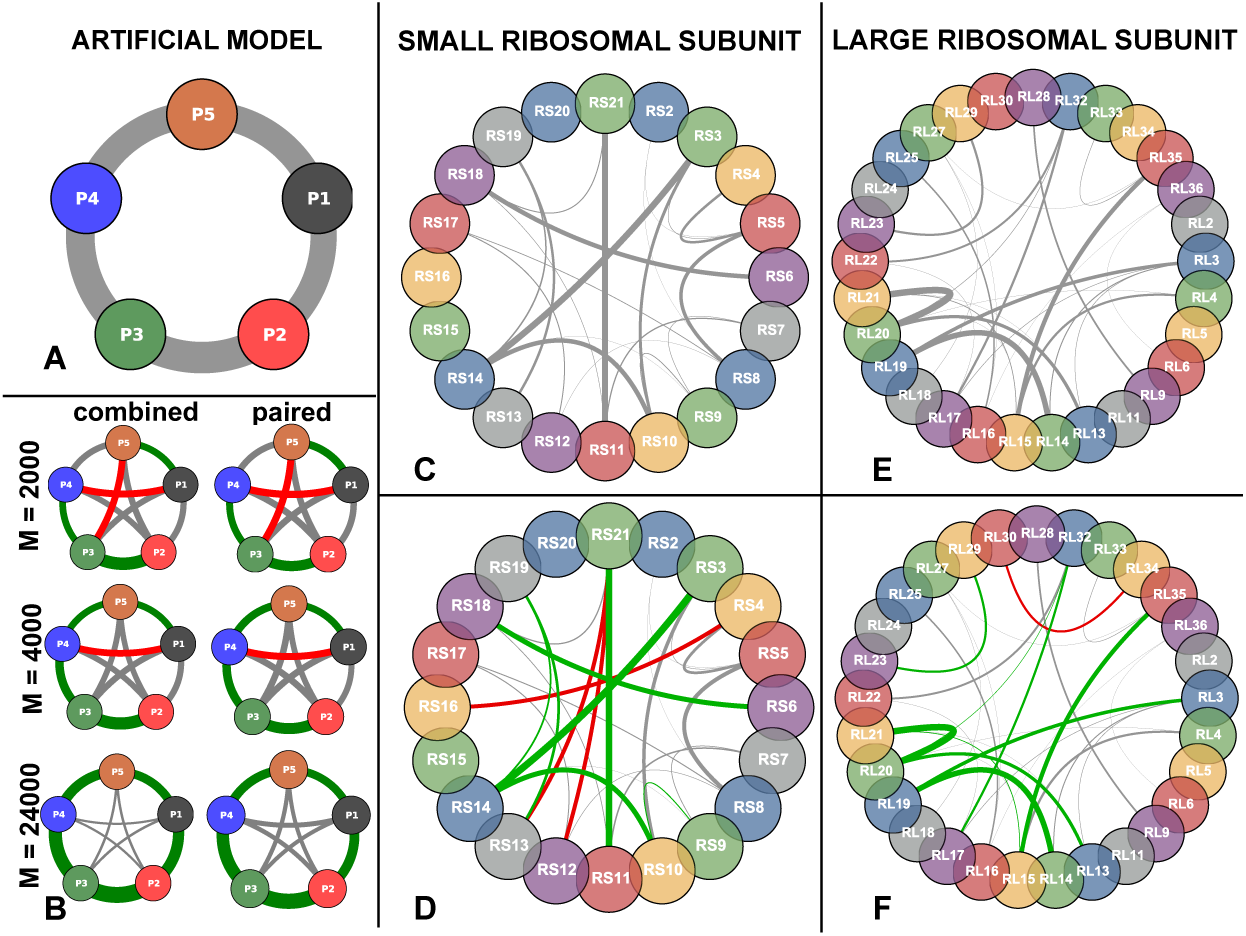
Residue-residue structure of both artificial and ribosomal complex. **A** Architecture of the *artificial* protein complex. Arcs width are proportional to the number of inter-protein residue contacts. **B** Inferred PPI network for both *paired* and *combined* strategy for different number *M* of sequences generated from the artificial model. Green arcs are true positives, red false positives, gray low-ranking predictions. Arc widths are proportional to the inter-protein interaction score. **C** SRU architecture (same color code as A). **D** Inferred PPI network (same color code as B). **E** Same as C for LRU. **F** Same as D for LRU. Arc width in panels C-F is provided by the number of inter-protein contacts, as a measure of interface size. It becomes obvious that mainly large interfaces are recognized by our approach.

To assess our capability to infer the PPI network of panel A from such data, we adopted two different strategies which we called *combined* and *paired* in panel B of Fig. 3. The *combined* strategy uses plmDCA on the full-length alignments of length 265 and models the interaction between all proteins pairs *simultaneously*. Given that in this artificial setting we use the same model to generate the data as to analyze it, the approach is guaranteed to infer the model correctly for a large number of analyzed sequences and therefore to assign a higher interaction score to any interacting protein pair than to any non-interacting pair.

To assess the coupling strength between two proteins, we average the four strongest residue coupling strengths between them. This leads to a score oriented toward the strongest signal while also reducing noise by averaging. In panel B of Fig. 3 we show the results for MSA sizes *M* = 2000, 4000, 24, 000 while intermediate values are reported in Figure S6 in S1 Text. The two lower figures - *M* = 2000, 4000 - represent the lower and upper bound of what we can currently obtain from databases for the proteins analyzed by us. The largest value *M* = 24, 000 is what we expect to be available in a few years from now, seen the explosive growth of sequence databases. The thickness of each link in Fig. 3 is proportional to the inferred inter-protein interaction score. The five strongest links are colored in green when they correspond to actual PPI according to panel A, and in red when they correspond to non-interacting pairs. For increasing sample size the predictions become more consistent and for *M* = 24, 000 any interacting protein-pair has a higher interaction score than any non-interacting pair.

Due to the running time of plmDCA only alignments for sequences of total length *L* ≾ 1000 can be analyzed. This is exceeded already by the sum of the lengths of the proteins of the small ribosomal subunit. Additionally, creating a combined multiple sequence alignment for more than two proteins would lead to very low sequence numbers due to the necessary matching (see *Materials and Methods*). Therefore, using the combined strategy is not generally applicable. In the *paired* strategy we therefore analyze each pair of proteins separately. This means that plmDCA is applied to all 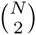 protein-pair alignments *D_ab_*, 1 ≤ *a < b ≤ N*. In panel B of Fig. 3 we find that the paired strategy is also able to detect the correct PPI network for large enough *M*. We observe, however, that the performance of the paired strategy is slightly worse. Couplings between non-interacting proteins are estimated significantly larger than using the combined strategy for large *M*. Even in the limit *M → ∞* we do not expect these links to disappear: Correlations between, e.g., *P*_1_ and *P*_3_ are generated via the paths 1 – 2 – 3 and 1 – 5 – 4 – 3, but in the paired strategy these correlations have to be modeled by direct couplings between *P*_1_ and *P*_3_ since the real direct coupling paths are not contained in the data.

After having answered the *’who-with-whom’* question for the artificial protein network, we address the *’how’* question of finding inter-protein contact pairs. Fig. 4 panel A displays individual residue contact pairs within and between proteins in the artificial complex. Panel B shows the 10 strongest intra-protein couplings for each protein and the 10 strongest inter-protein couplings inferred by plmDCA (*M* = 4000, combined strategy). Green links correspond to contact pairs and red links to non-contact pairs. We see that the intra-protein prediction is perfect, whereas a few errors appear for inter-protein predictions in agreement with the results of Fig. 3.

### The PPI network of bacterial ribosomes

As a more realistic test we apply the method to the bacterial large and small ribosomal subunits (LRU, SRU). To define contacts and protein interaction partners we used high-resolution crystal structures with PDB-IDs 2z4k (SRU) and 2z4l (LRU) [4]. The contact network is summarized by the contact maps in Fig. 1. The ribosomal RNA is ignored in our analysis.

Panels C, E of Fig. 3 display the architectures of both SRU and LRU. The SRU (LRU) complex consists of 20 (29) proteins of lengths 51–218 (38–271); 21 (29) out of 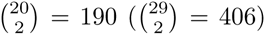 pairs are in contact. The interfaces contain between 3–209 (1–229) residue pairs. The width of the inter-protein links in the PPI network Fig. 3 in panels C, E are proportional to these numbers. The number of contacts within the individual proteins ranges from 297 to 2337 (303–2687). Globally, there are 22644 (30555) intra-protein and 1401 (1,439) inter-protein contacts, so the contacts relevant for our study comprise only 5.8% (4.5%) of all contacts.

Fig. 3 panel D shows the inferred SRU PPI architecture. As expected, the biological case is harder than the artificial case where the data are independently and identically distributed according to the generating model. Even though the histograms of the inferred interaction scores for both cases are very similar (see Figure S2 in S1 Text, biological data are expected to show non-functional correlations due to the effect of phylogeny or sequencing efforts which are biased to model species and known pathogens. Nonetheless, among the top ten predicted interacting protein pairs the method makes only three errors (true-positive rate 70% as compared to 21/190 ≃ 11% true PPI between all protein pairs, with an overall area under ROC curve (AUC) of 0.69 (see Fig. 7). The method spots correctly the pairs with larger interaction surfaces whereas the small ones are lost. Two of the false-positive (FP) predictions include protein RS21, which has the smallest paired alignments with other proteins (*M* between 1468 and 1931). Also the third FP, corresponding to the pair RS4-RS18, is probably due to a small MSA with *M* = 2064. At the same time, the interaction of RS21 with RS11, which is one of the largest interfaces (199 contacts), is still detected despite the low *M* = 1729. The same procedure for the LRU (406 protein pairs) performs even better: 9 out of the 10 first PPI predictions are correct (see Fig. 3 panel F), and the AUC is 0.81.

The results on the residue scale for both SRU and LRU are depicted in panels D and F of Fig. 4. Shown are the first 20 intra-protein residue contact predictions for each protein (excluding contacts with linear sequence separations below 5 to concentrate on non-trivial predictions) and the first 20 inter-protein residue contact predictions. In the SRU case of panel D for example, the results are qualitatively similar to the artificial case, albeit with a slightly reduced true-positive rate of 60% among the first 20 inter-protein residue contact predictions (compared to the ratio of 1401 actual inter-protein residue contacts and 2,403,992 possible inter-protein residue contacts, i. e., 0.058%). Again 3 out of the 8 false positives are related to RS21, which due to the smaller MSA size is also the only one having a considerable false-positive rate in the intra-protein residue contact prediction. About 95% of the displayed 400 highest intraprotein residue contacts are actually contacts (see Figure 1 in S1 Text). Analogous considerations with a somewhat larger accuracy (85%) hold for LRU as displayed in Fig. 4 panel F.

### The PPI network of the tryptophan biosynthetic pathway

As a distinct test case for our methodology we analyzed the 7 enzymes (TrpA, B,C,D,E,F,G) that comprise the well characterized tryptophan biosynthesis pathway. In contrast to the ribosomal proteins, these enzymes are only conditionally essential in the absence of environmental tryptophan and their genes are only expressed under deplete tryptophan conditions. In this particular system, only two protein-protein interactions are known and resolved structurally: TrpA-TrpB (PDB-ID 1k7f [40]) and TrpG-TrpE (PDB-ID 1qdl [27]). Whereas the TrpG-TrpE pair catalyzes a single step in the pathway and their interaction is thus essential for correct functioning, the TrpA-TrpB pair catalyzes the last two steps in tryptophan biosynthesis. Both enzymes function in isolation but their interactions are known to increase substrate affinity and reaction velocity by up to two orders of magnitude. All other proteins catalyze individual reactions, but one might speculate that the efficiency of the pathway could benefit from co-localization of enzymes involved in subsequent reactions. Interestingly, the Pfam database [15] reports that in many species pairs of genes in the operon appear to be fused, suggesting that some of the fused pairs are actually PPI candidates. An example is the TrpCF protein, which is fused in *Escherichia coli* and related species (but not in the majority of species).

After applying our method to all 21 protein pairs we find elevated interaction scores only for TrpA-TrpB and TrpE-TrpG, which are the only known interacting pairs (see Fig. 5 and Table S10 in S1 Text for the interaction scores of all pairs). Those two pairs have interaction scores of 0.375 and 0.295, while the other pairs are distributed between 0.071 and 0.167. Even though we do not define a significance threshold for prediction (see Section *A global view*), these two pairs would be discernible as interesting candidates even if we did not have the 3D structures.

We speculate therefore that the fusions in many species do not imply strong interprotein co-evolution. To further investigate this aspect, we took a closer look at the protein pair TrpC-TrpF. For this protein pair, a high resolution structure of a fused version exists (PDB-ID 1pii [41]). We ran our algorithm on the complete multiple sequence alignment, the multiple sequence alignment with fused sequence pairs removed and only on the fused sequences. In none of these cases did we observe a statistically significant interaction score or a statistically significant prediction of inter-protein contacts present in the structure of the fused protein.

Our results are corroborated by the finding that all scores measuring the co-evolution between a ribosomal protein and an enzyme from the tryptophan synthesis pathway are small (see the following subsection). No indication for an interaction between the two systems is found, as to be expected from the disjoint functions of the two systems.

### A global view

It is interesting to assemble a larger-scale system out of the three systems (SRU, LRU, Trp). To this end, we created all possible pairings between the proteins used in the present study (SRU vs. RU, SRU vs. Trp, LRU vs Trp, SRU vs SRU, LRU vs. LRU, and Trp vs. Trp). This leads to a total of 1540 pairs, out of which only 49 pairs are known to interact (which we defined as true positives). We present the findings in Fig. 7 and in Figs. S7-S9 in S1 Text. Fig. S7 in S1 Text shows the true-negative rate, which is the fraction of true negatives in the indicated number of predictions with the *lowest* interaction scores. As it can be seen our scoring produces a false negative just after 420 true negatives. Figs. 7 and S8 show true positive rates for the complete system and the individual systems. We also show true positive rates for alternative ways to calculate the interaction score between protein pairs, i.e. a different number of inter-protein residue-residue interaction scores to average. We notice that in the complete system, the performance is similar to the performance in individual systems. All of the 10 highest-scoring protein pairs are known to interact, and 75% of the first 20 protein-pairs. After these first 20 pairs, the true positive rate drops to around 45% in the first 40 predictions. This is analogous to the case of protein contact prediction, where methods based on the same model are able to extract a number of high confidence contacts but see a large drop in performance afterwards [31]. The area under the AUC for the whole system is 0.83 (see Fig. 7). This is stable when averaging different numbers of residue-contact scores to arrive at a protein-protein interaction score, but the performance seems to worsen when using more than 6. This is probably because only a few inter-protein residue contacts have a large score and averaging over too many only adds noise. It can also be seen that averaging over 4 performs very well in the small ribosomal subunit, which is why we have chosen this value for the large part of the analysis. On the larger-scale system, though, any number between 1 and 6 performs almost identically.

A further question is whether it is possible to define a threshold allowing to reliably discriminate between interacting and non interacting pairs in terms of the interaction score. Figure S9 in S1 Text shows two normalized histograms of the interaction scores. The rightmost tail of the interacting pairs distribution is well separated from the rightmost tail of non-interacting one, but the highest scores of non-interacting pairs are strongly overlapping with the lowest scores of the interacting ones. The situation is therefore analogous to what is observed in the case of the inference of contacts within single protein families [2, 12, 31, 38], where the same technique is known to produce relatively few high confidence contacts in the topmost scoring residue pairs. To conclude, while high scores seem to reliably predict interacting pairs, and low scores non-interacting pairs, there is a large gray zone prohibiting a clear discrimination between interacting and non-interacting pairs.

**Figure 4:**
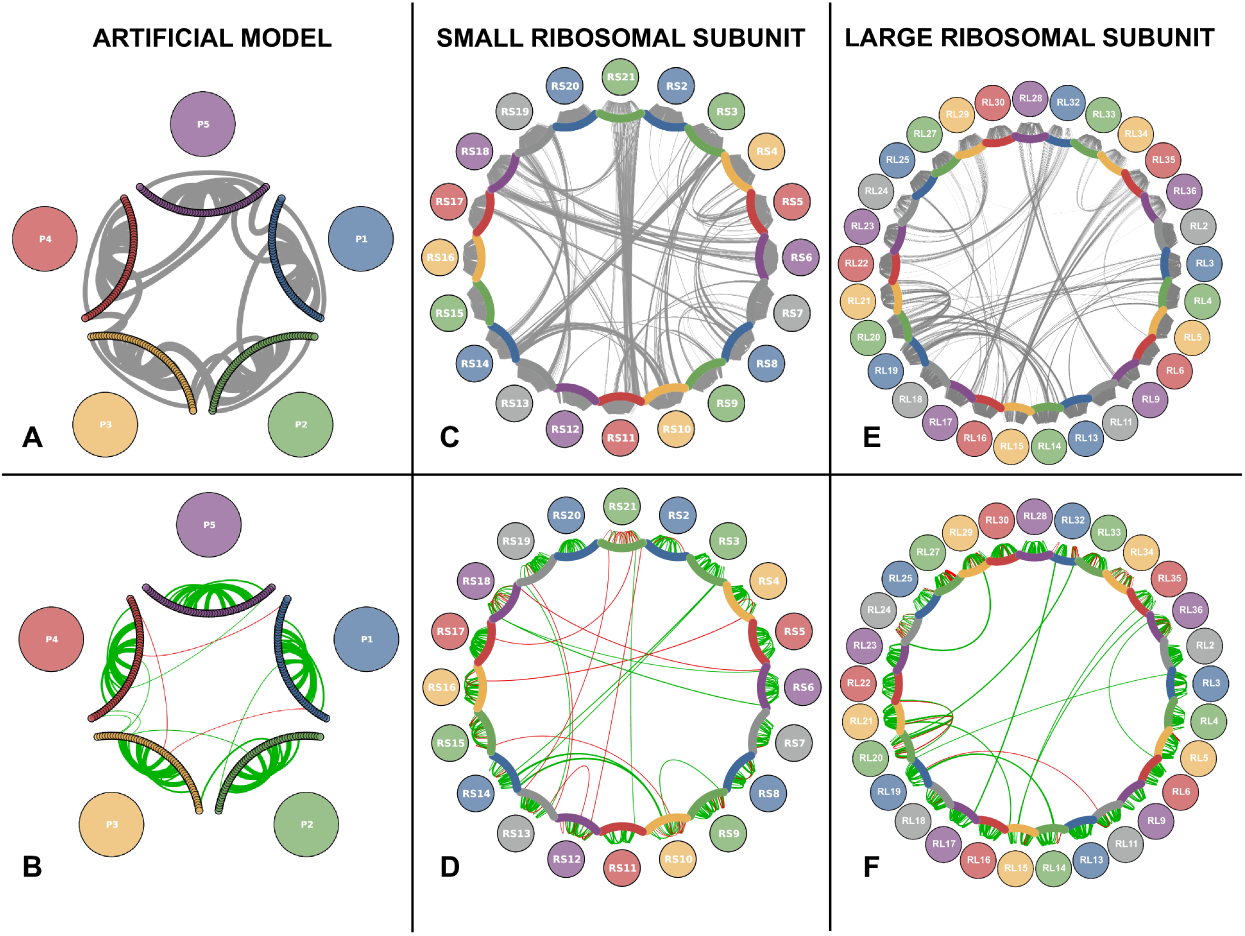
Architecture and inferred protein-protein interaction network of the artificial protein complex. **A** Residue-residue interaction structure of the generating model for the artificial data. Colored arcs represent the protein chain. Non-zero couplings in the coupling matrix of the generating model are represented as curves between the nodes. The width of the curves is proportional to the interaction score. Only the 10 strongest intra/interprotein scores are shown. **B** Same as **A**, but based on the inferred couplings. Green arcs are true positives, red false positives. Note that not all green arcs have a corresponding arc in **A** due to our choice to display only the 10 strongest couplings, which not always correspond to the strongest score. **C** Same as **A** for SRU. All links represent a contact in the PDB structure and have equal width. **D** Same as **B** for SRU. **E** Same as **C** for LRU. All links represent a contact in the PDB structure and have equal width. **F** Same as D for LRU.

**Figure 5:**
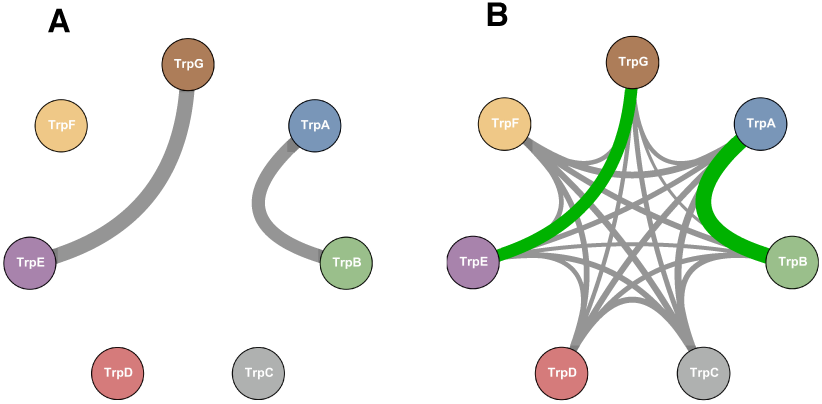
Tryptophan biosynthesis pathway. A Architecture of the known proteinprotein interaction among the 7 enzymes which are coded in the Trp operon. The widths of the arcs are proportional to the number of inter-protein residues (which in this case is almost equal for the two interacting pairs). **B** Inferred PPI network, here the width of the arcs is proportional to the interaction score. Green arc correspond to the protein pairs for which a known structure exist.

**Figure 6:**
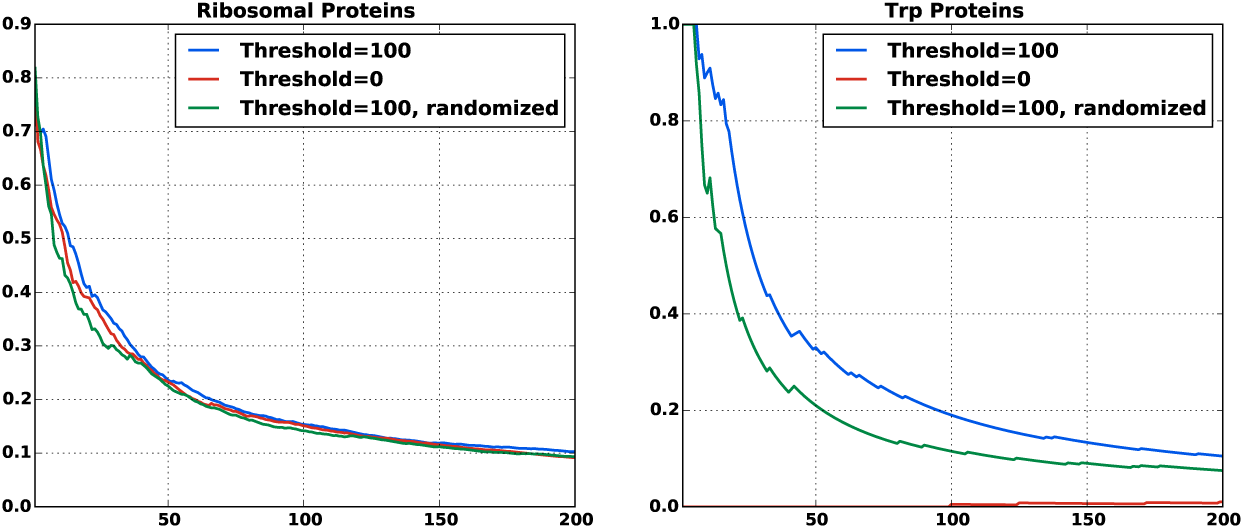
Efficacy of the different matching procedures. True-positive rates for inter-protein residue contact prediction for different matching procedures. Shown are means for all protein pairs that have at least 100 residue pairs in contact. The ribosomal and the trp proteins were tested independently. The red curves correspond to a matching including only protein sequences without paralogs inside the same species (”matching by uniqueness in genome”). The low performance of this approach on Trp proteins is due to a very low number of species without homologs, which leads to very small matched alignments. The blue curves show the results for our matching procedure as described in the text. The green curves correspond to alignments that have been obtained by first applying our matching procedure and then randomizing the matching within individual species. The definition of “contact” was the same as used above (a distance of less than 8.0Abetween two heavy atom in the residues).

## Conclusions

To conclude, we have shown that DCA performs excellently in the systems tested when used to predict protein-protein interaction partners. In the small and large ribosomal subunit our tests resulted in a true positive rate of 70% and 90% in the first 10 predictions (AUC of 0.69 and 0.81) while in the trp operon the two largest interaction scores corresponded to the only two interactions experimentally known (AUC 1). The performance is summarized in Fig. 7. The Figure shows both the high quality of the first predictions, but also a drop in performance after a fraction of all interacting pairs (about 40% in our test case). This is analogous to the case of protein contact prediction by DCA and related methods, where the performance drops after a limited number of high-confidence predictions [31]. In the same context and with the same caveat, an excellent performance in predicting inter-protein contacts on the residue level has been shown. The artificial data have shown that the performance of our approach depends crucially on the size of the alignments. Only for very large MSA (*M* = 24, 000 sequences in our data) a perfect inference of the artificial PPI network was achieved. MSA for real proteins pairs are typically much smaller. Even for pairs of ribosomal proteins, which exist in all bacterial genomes, only about 1500–3200 sequence pairs could be recovered. This places these data towards the lower detection threshold of PPI. We therefore expect the performance of the presented approach to improve in the near future thanks to the ongoing sequencing efforts (the number of sequence entries in Uniprot [8] has been growing from about 10 millions in 2010 to 90 millions in early 2015) and improved inference schemes. The strong performance of the same algorithm on different and dissimilar systems naturally prompts us to expect that the approach can be used to detect interactions experimentally unknown so far. In fact, if we trust our results on the trp operon we can already draw some speculative biological inferences. While there are many high-resolution structures of the ribosome available, one might have expected that in the trp operon there could be more transient previously unreported interactions in the tryptophan biosynthesis pathway beyond the two interactions that have been structurally characterized. As mentioned, various enzyme fusions can be observed in the databases, suggesting that there is an evolutionary benefit to co-localizing the enzymes of the pathway in the cell. An obvious benefit of such co-localization would be that the pathway intermediates do not have to diffuse throughout the cell from one enzyme component to the next. In the tryptophan biosynthesis pathway in particular, there are numerous phosphorylated intermediates that need to be protected from unspecific cellular phosphatase activity. Organizing the enzymes in the pathway in a multi-protein complex would seem like an efficient way to protect the intermediates from decay. However, our data indicate that the only statistically relevant co-evolutionary signals that can be observed are restricted to the known strong interactions between TrpA with TrpB and TrpE with TrpG. This could be interpreted in a number of ways: *(i)* The most obvious explanation is that there are no additional protein-protein interactions beyond those that are known and that no multi-enzyme complex exists for the tryptophan biosynthesis pathway. Alternatively *(ii)* it seems plausible that there are numerous structural solutions to form a tryptophan biosynthesis complex and that there is no dominant structure from which a co-evolutionary pattern can be observed in the sequence databases. Lastly *(iii)* it is not out of question that the enzymes of the pathway do not directly form a complex but that they are jointly interacting with an unidentified scaffold component. Of course we cannot exclude that our method is not able to capture other potentially present interactions.

**Figure 7:**
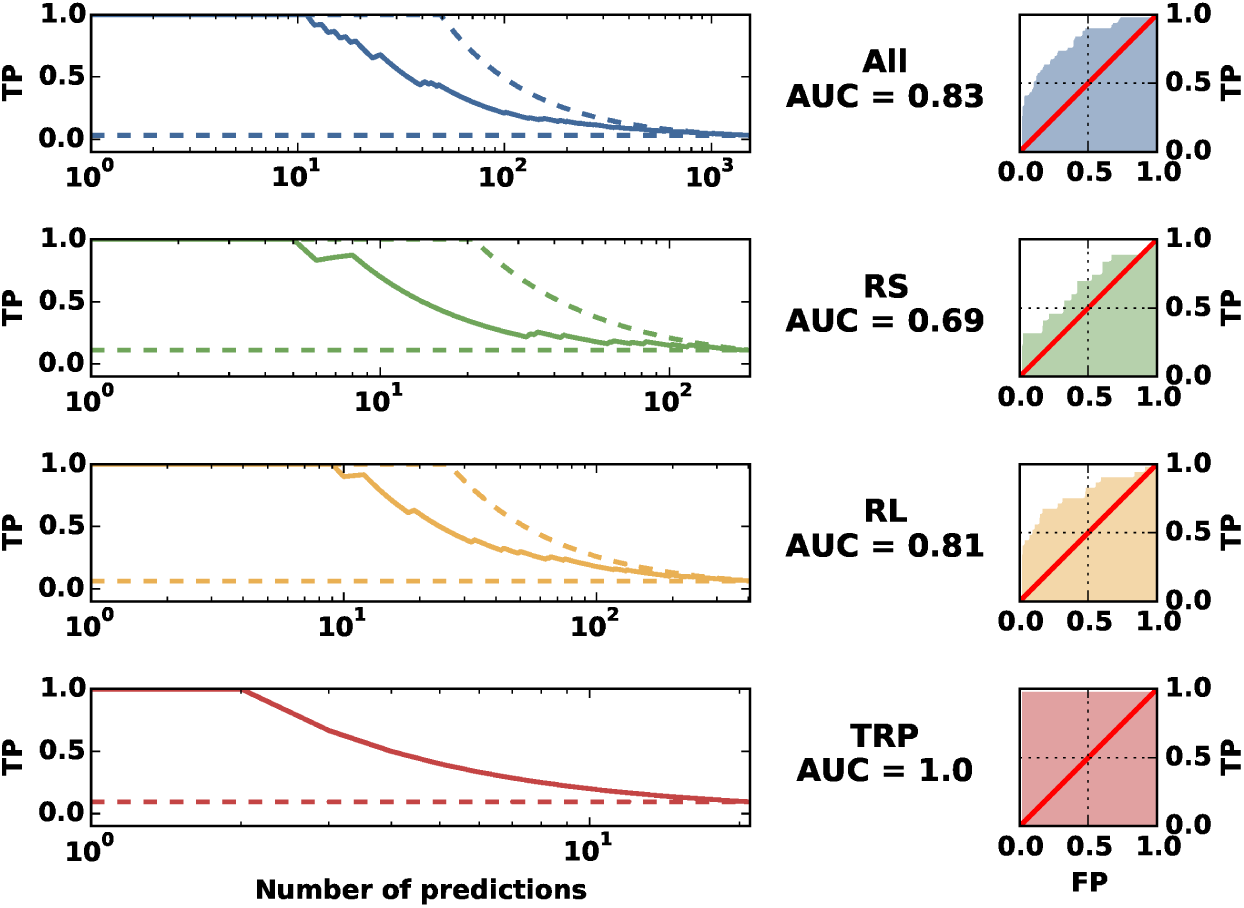
Performance Summary. The plots illustrate the performance in predicting protein interaction partners. The left panels show the fraction of true positives among the first *n* PPI predictions, with *n* being the number indicated on the horizontal axis (solid lines). The dashed lines show the best possible (upper dashed line) and the mean of a random prediction (lower dashed line). The right panels show ROC-curves, which indicate the dependence of the true-positive predictions (TP/P) from the false positive predictions (FP/N). The area under the curve (AUC) is a global global measure for the prediction quality; it is 1/2 for a random, and 1 for a perfect prediction. A protein pair is identified as an interacting (true positive) pair, if at least one PDB structure with at least one inter-protein contacts exists.

From a methodological point of view, one possible algorithmic improvement is creating better MSAs for protein pairs. The vast majority of protein families show genomic amplification within species. This raises the issue of which sequence in one MSA should be matched with which sequence in the other MSA when concatenating the two MSAs, as shown in Fig. 2. In the absence of prior knowledge and as long as only prokaryotes are concerned, we showed that it is possible to use the simple criterion of *matching by genomic proximity*. This criterion is based on the observation that two sequences are more likely to interact if they are genomically co-localized. Our results have shown that in the case of the ribosomal network better inference results can be obtained by using this matching criterion than by using a random matching or using a conservative matching taking only species with a single sequence in both MSAs into account, cf. Fig. 6. However, we found it beneficial for the predictive performance to introduce a threshold distance above which we simply discarded candidate sequences. This is not based on biological principles.

We believe that our *naive* matching strategy can be improved substantially. Even if closeness of sequence pairs on the genome is a good proxy for interaction in some cases, for example if they belong to the same operon, excluding all distal pairs is a very crude criterion. This criterion is known to be erroneous in many cases, for example in the bacterial two component signal transduction system [6, 7, 35]. It would therefore be interesting to include the matching into the inference procedure itself, *e.g*. to find a matching that maximizes the inter-protein sequence covariation, cf. [6] for a related idea. However, for highly amplified protein families this leads to a computationally hard optimization task. Simple implementations get stuck in local minima and do not lead to improvements over the simple and straight-forward scheme proposed here.

## Acknowledgments

CF and AP are supported from the EU Marie Curie Training Network NETADIS, (FP7 Grant 290038). HS was supported by grants GM106085 and GM019416 from the US National Institute of General Medical Sciences, National Institutes of Health. MW was supported by the Agence Nationale de la Recherche project COEVSTAT (ANR-13-BS04–0012–01). CF and AP acknowledge Riccardo Zecchina and Carlo Baldassi for many interesting discussions.

## Supporting Informations

**S1 Text**

## 1 Multiple Sequence Alignments

### 1.1 Multiple Sequence Alignments

The data we use are multiple sequence alignments (MSA). Each such MSA is a rectangular matrix, with entries coming from a 21-letter alphabet containing the 20 standard amino acids and a gap symbol “−”. In the following we denote this alignment by a matrix

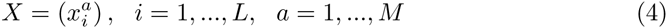

with *L* being the number of residues of each MSA row, i.e., the number of residues in each considered protein, and *M* the number of MSA rows, i.e., the number of proteins collected in the alignment. For simplicity of notation we assume that the 21 amino acids are translated into consecutive numbers 1,…,21.

### 1.2 Alignment Generation

For all proteins of the small ribosomal subunit (SRU) and the large ribosomal subunit (LRU) the sequence names were extracted from the corresponding PFAM alignments [15]. Using these names, the following procedure was used to create the alignments for the single proteins:

1. Extract sequences corresponding to names from Uniprot [8]
2. Run MAFFT [26] on them using `mafft --anysymbol --auto`
3. Remove columns from the alignment that contain more than 80% gaps
4. Create an Hidden Markov Model (HMM) using `hmmbuild` from the hmmer suite [17]
5. Search Uniprot using `hmmsearch` [17]
6. Remove inserts
7. If there exist in one species two or more sequences that are more than 95% identical, remove all but one.

The number of sequences for the single files can be found in Table 1

The alignments for the proteins of the Trp Operon where constructed analogously with some modifications to ensure that only full-length sequences where extracted. Also, we chose the `linsi` program of the MAFFT package to create the initial MSAs. The number of sequences for the Trp alignments can be found in Table 2.

**Table 1:** Alignment sizes (M) and lengths (L) for proteins of the small (RSXX) and large (RLXX) ribosomal subunit. (P) indicates the average number of paralogs per species and (S) the standard deviation of this number.

|  | L | M | P | S |
| --- | --- | --- | --- | --- |
| RS2 | 219 | 6053 | 1.743 | 5.978 |
| RS3 | 216 | 6235 | 1.716 | 7.761 |
| RS4 | 171 | 8522 | 2.175 | 11.305 |
| RS5 | 164 | 5075 | 1.678 | 5.845 |
| RS6 | 105 | 4132 | 1.563 | 6.630 |
| RS7 | 147 | 5733 | 1.595 | 4.962 |
| RS8 | 127 | 5761 | 1.700 | 5.992 |
| RS9 | 127 | 4983 | 1.663 | 5.917 |
| RS10 | 100 | 4560 | 1.511 | 4.232 |
| RS11 | 120 | 5136 | 1.520 | 4.019 |
| RS12 | 124 | 5607 | 1.581 | 4.036 |
| RS13 | 116 | 5729 | 1.856 | 5.763 |
| RS14 | 96 | 5555 | 1.689 | 4.780 |
| RS15 | 89 | 5361 | 1.646 | 6.036 |
| RS16 | 83 | 4463 | 1.507 | 5.851 |
| RS17 | 82 | 4774 | 1.616 | 5.481 |
| RS18 | 73 | 4512 | 1.483 | 4.879 |
| RS19 | 89 | 5364 | 1.537 | 4.700 |
| RS20 | 88 | 3848 | 1.676 | 7.460 |
| RS21 | 65 | 3209 | 1.456 | 4.188 |
| RL3 | 205 | 6077 | 2.025 | 6.522 |
| RL4 | 198 | 5671 | 1.906 | 6.810 |
| RL5 | 177 | 5032 | 1.636 | 6.245 |
| RL6 | 178 | 5308 | 1.765 | 6.894 |
| RL9 | 149 | 4199 | 1.698 | 7.621 |
| RL11 | 141 | 5027 | 1.683 | 5.517 |
| RL13 | 147 | 5091 | 1.717 | 6.458 |
| RL14 | 120 | 5145 | 1.528 | 4.358 |
| RL15 | 140 | 5926 | 1.964 | 6.754 |
| RL16 | 133 | 5673 | 1.604 | 4.904 |
| RL17 | 121 | 4345 | 1.612 | 7.637 |
| RL18 | 111 | 4961 | 1.674 | 6.570 |
| RL19 | 116 | 4079 | 1.511 | 6.454 |
| RL20 | 119 | 4476 | 1.554 | 5.864 |
| RL21 | 102 | 4123 | 1.551 | 6.486 |
| RL22 | 108 | 6378 | 1.918 | 5.790 |
| RL23 | 87 | 5632 | 1.711 | 6.292 |
| RL24 | 99 | 9062 | 3.073 | 12.820 |
| RL25 | 186 | 3272 | 1.680 | 6.109 |
| RL27 | 89 | 3989 | 1.486 | 5.419 |
| RL28 | 74 | 4051 | 1.584 | 5.694 |
| RL29 | 66 | 4456 | 1.540 | 6.024 |
| RL30 | 60 | 4356 | 1.671 | 5.313 |
| RL32 | 60 | 4206 | 1.463 | 4.997 |
| RL33 | 49 | 4604 | 1.678 | 4.943 |
| RL34 | 45 | 3195 | 1.346 | 4.280 |
| RL35 | 65 | 3691 | 1.502 | 5.889 |
| RL36 | 38 | 3779 | 1.408 | 3.103 |

**Table 2:** Alignment sizes (M) and lengths (L) for proteins of the Trp Operon. (P) indicates the average number of paralogs per species and (S) the standard deviation of this number.

|  | L | M | P | S |
| --- | --- | --- | --- | --- |
| TrpA | 259 | 10220 | 4.457 | 32.604 |
| TrpB | 399 | 46557 | 16.992 | 145.826 |
| TrpC | 254 | 10323 | 4.536 | 39.868 |
| TrpD | 337 | 17582 | 7.130 | 59.693 |
| TrpE | 460 | 28173 | 11.749 | 124.933 |
| TrpF | 197 | 8713 | 4.122 | 32.400 |
| TrpG | 192 | 78265 | 24.713 | 187.331 |

### 1.3 Internal Sensitivity Plots

As an assessment of quality for the alignments, sensitivity plots using the pdb files `2Z4K` and `2Z4L` were made. Figure 8 shows results for contact predictions based on the GaussDCA [2] and plmDCA alghorithm [13].

**Figure 8:**
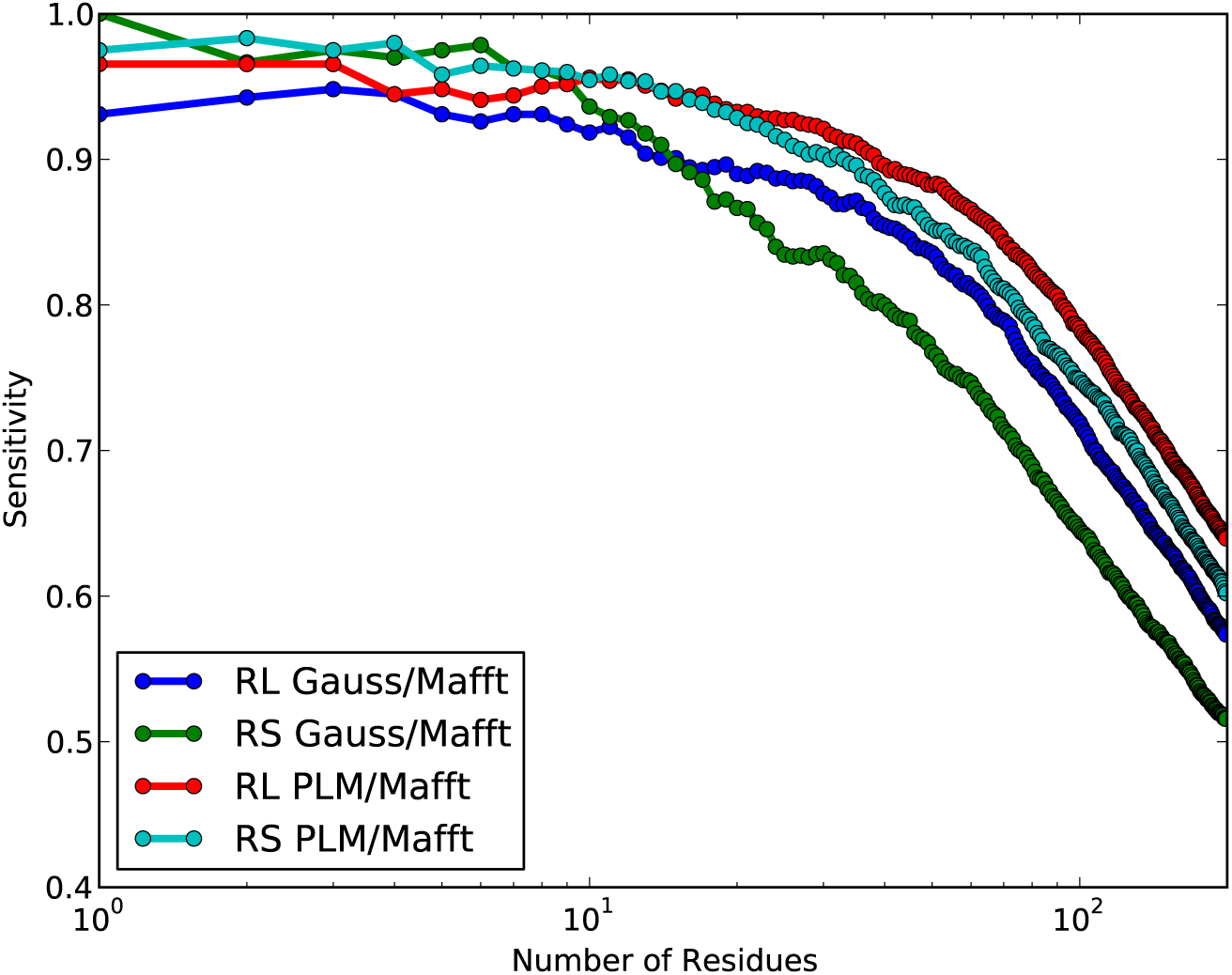
Intra-Protein Sensitivity Plots. On the alignments for the single ribosomal proteins the plmDCA algorithm was run and an ordered list of residue pairs obtained. For every number *n* on the abscissae the fraction of the number of true positives (the sensitivity) in the first *n* pairs on this list was calculated for every protein. The plot shows the mean of these values for the Gaussian algorithm of [2] and the plmDCA algorithm run on the proteins of the large and small ribosomal subunit.

## 2 Matching Procedure

### 2.1 Pipeline for Matching

The problem of generating a concatenated alignment from two MSAs of two different protein families (say MSA_1_ and MSA_2_) is to decide which sequence from the first alignment should be concatenated to which sequence from the other alignment. This means to find for any protein 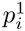 in MSA_1_ a matching partner 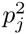 in MSA_2_ belonging to the same species. The problem is trivially solved in the case when no paralogs are present and each species has one and only one sequence in each individual MSA. In this case we can simply concatenate these two sequences (we term this case *matching by uniqueness*). The problem is that species often have several paralogs. In this case, given that we would like to observe a co-evolutionary signal between protein interaction partners, one would like to match sequences of proteins that are (possibly) interacting.

As long as Prokaryotes are concerned, it turns out empirically that proteins are more likely to interact if their genes are *co-localized* on the DNA [6, 38]. This suggests to try to match proteins that are close on the genome when creating a concatenated MSA.

As a proxy to the genomic distance we use a *distance* between Uniprot accession numbers (UAN). This UAN consists of a 6 digit alphanumeric sequence for every sequence and can be extracted from the sequence annotation, e.g. the `”D8UHT6”` part of the sequence annotation `”D8UHT6_PANSA”`.

We define the distance between UANs as follows: Different positions in the UAN can take on different values, some only numeric (0–9) and some alphanumeric values (0–9,A-Z). We define for every position *i* ∈ 1… 6 the number *B_i_* as the number of different values position *i* can take, i.e. *B_i_* = 10 for the numeric positions and *B_i_* = 36 for the alphanumeric positions.

We further map the possible single position values in the UAN to the natural numbers in ascending order, i.e. we assign to the numeric symbols 0–9 the natural numbers 0 – 9 and to the letters the natural numbers following 9 (so to A we assign 10, to B we assign 11 etc.). This leads for example for the the UAN `L9XG27` to the numeric sequence *A* = (21, 9, 33, 16, 2, 7).

Now we can define a unique number *N* for any UAN that has been mapped to the sequence of natural numbers *A_i_* as

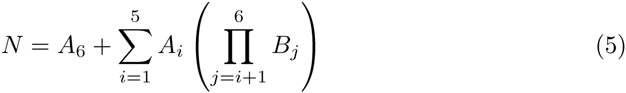

The distance between two UANs that have been mapped to the numbers *N*_1_ and *N*_2_ can now be defined as

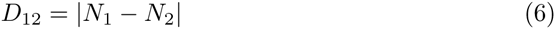

This procedure induces a distance *D_ij_* for any sequence *p_i_* ∈ MSA_1_ and *p_j_* ∈ MSA_2_, where both *p_i_*, *p_j_* belong to the same species. In this way we define a complete weighted bipartite graph, and the problem of finding the proper pairing can thus be translated into a minimum weighted bipartite matching problem. This problem can be readily solved using a standard linear programming techniques. Finally we discard from the optimal solution sequence pairs whose distance is above a given threshold of 100 (manually optimized on the small ribosomal subunit). In the cases we analyzed, such a threshold moderately increases the quality of the prediction of interaction partners.

## 3 Inference technique

As a simple but meaningful statistical model, we consider a pairwise generalized 21 states (to mimic the 20 amino acids + 1 insert symbol alphabet of MSAs) Potts model with the following Hamiltonian

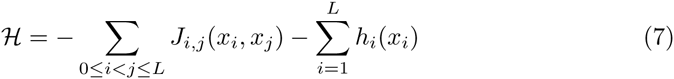

We can now assume to have a dataset *D* = {*x*^1^,…,*x^M^*}, where *x* represents one sequence, either artificially generated, or extracted using the bioinformatic pipeline discussed above. Notice that if the sequences *x* are concatenations of two sequences (*x, x*^′^), the sums in Equation 7 can be split into three parts: One in which appear only sites in *x*, one in which appear only sites in *x*′and one interaction part with *J_ij_* for which *i* is in *x* and *j* in *x′*. By labeling the first part *H*(*x*), the second *H*′(*x*′) and the third H*^int^*(*x, x*′) one arrives at the representation referred to in the main text. Given that the representations are mathematically equivalent, we will here in supplemental information treat the sequence as one simple sequence *x*.

The inference proceeds by assuming as a working hypothesis that the dataset *D* is composed by configuration sampled uniformly from the equilibrium Boltzmann-Gibbs distribution 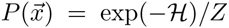 (as an inference process, we are free to consider *T* = *β* = 1). We are now ready to use *D* to infer the topology of the network. To do so – as discussed in the main text – in the last years different maximum-likelihood techniques have been proposed [1, 12, 23, 29, 31, 39]. So far the most promising in terms of accuracy seems to be the pseudo-likelihood maximization introduced in [12] where from the previously defined Boltzmann-Gibbs measure we consider the following conditional probability distribution:

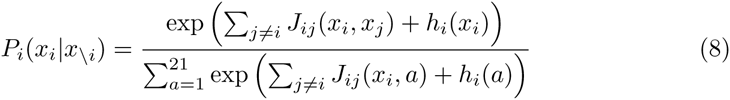

Given a data set *D* we can thus maximize the conditional likelihood by maximizing

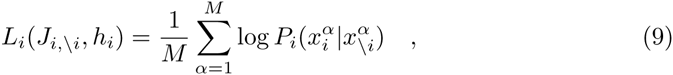

as a function of J*_i_,_\i_, h_i_*. As customary in many maximum-likelihood inference techniques, we add to the maximization an 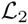 regularization term, so that eventually the extremization procedure turns out to be:

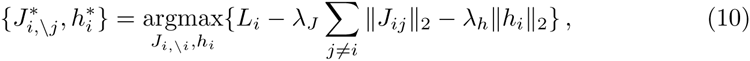

with 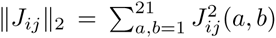, and 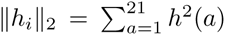. We refer to the original paper [12] for the details of the implementation. We only mention that beside the original MATLAB [30] implementation available at http://plmdca.csc.kth.se/, we developed an efficient implementation of the pseudo-likelihood implementation in a new open-source language called Julia [3]. The package can be downloaded at https://github.com/pagnani/PlmDCA.

## 4 Ribosomal Protein Interaction Partner Prediction

Using the ribosomal alignments as described in Section 1 and the matching as described in Section 2, concatenated alignments for the ribosomal proteins (small and large ribosomal subunit independently) were created. Tables 4 and 3 show the resulting alignment sizes for the SRU and Tables 6 and 5 for the LRU.

The creation of the alignments for the Trp Proteins was analagous and the resulting alignment sizes can be found in Table 7.

As discussed in the main text, in principle one would be interested in a MSA in which a sequence is a concatenation of sequences from all proteins families in the complex at once. A comparative glance at Tables 5 and 1 shows that in the matching procedure described above a lot of sequences have to be discarded for not having a suitable matching partner. This leads to a reduction of the predictive power of the method. It is expected that extending the matching procedure to more than two proteins would lead to very low sequence numbers in the matched alignments and in turn reduce the predictive power of the method further. For this reason we only performed the concatenation of pairs of proteins.

**Table 3:** Matched Alignment Sizes for Small Ribosomal Subunit, at threshold 100

|  | RL2 | RL3 | RL4 | RL5 | RL6 | RL7 | RL8 | RL9 | RL10 | RL11 | RL12 | RL13 | RL14 | RL15 | RL16 | RL17 | RL18 | RL19 | RL20 | RL21 |
| --- | --- | --- | --- | --- | --- | --- | --- | --- | --- | --- | --- | --- | --- | --- | --- | --- | --- | --- | --- | --- |
| RL2 |  | 2914 | 2537 | 2458 | 2224 | 2825 | 2833 | 2491 | 2457 | 2839 | 2664 | 2342 | 2511 | 2748 | 2462 | 2373 | 2515 | 2842 | 2109 | 1740 |
| RL3 |  |  | 2947 | 2719 | 2430 | 3109 | 3223 | 2531 | 2680 | 3097 | 2922 | 2577 | 2992 | 2694 | 2645 | 2686 | 2659 | 3213 | 2123 | 1907 |
| RL4 |  |  |  | 2411 | 1837 | 2719 | 2812 | 2214 | 2314 | 2802 | 2528 | 2463 | 2522 | 2319 | 2064 | 2354 | 2182 | 2765 | 1774 | 1468 |
| RL5 |  |  |  |  | 2231 | 2613 | 2736 | 2508 | 2607 | 2623 | 2410 | 2532 | 2517 | 2381 | 2221 | 2699 | 2142 | 2657 | 2127 | 1743 |
| RL6 |  |  |  |  |  | 2206 | 2251 | 2216 | 2200 | 2204 | 2041 | 2117 | 1938 | 2169 | 2430 | 2226 | 2590 | 2263 | 2116 | 1931 |
| RL7 |  |  |  |  |  |  | 3001 | 2469 | 2580 | 2914 | 3172 | 2452 | 2753 | 2650 | 2414 | 2524 | 2483 | 2937 | 2089 | 1711 |
| RL8 |  |  |  |  |  |  |  | 2539 | 2782 | 3098 | 2831 | 2654 | 3004 | 2707 | 2494 | 3037 | 2497 | 3402 | 2114 | 1786 |
| RL9 |  |  |  |  |  |  |  |  | 2466 | 2564 | 2348 | 2400 | 2284 | 2383 | 2204 | 2469 | 2188 | 2489 | 2103 | 1755 |
| RL10 |  |  |  |  |  |  |  |  |  | 2579 | 2423 | 2460 | 2443 | 2378 | 2212 | 2711 | 2144 | 2784 | 2100 | 1734 |
| RL11 |  |  |  |  |  |  |  |  |  |  | 2810 | 2618 | 2849 | 2694 | 2417 | 2604 | 2497 | 3008 | 2083 | 1729 |
| RL12 |  |  |  |  |  |  |  |  |  |  |  | 2295 | 2646 | 2507 | 2224 | 2369 | 2303 | 2828 | 1925 | 1542 |
| RL13 |  |  |  |  |  |  |  |  |  |  |  |  | 2395 | 2188 | 2174 | 2502 | 2117 | 2564 | 2060 | 1712 |
| RL14 |  |  |  |  |  |  |  |  |  |  |  |  |  | 2420 | 2169 | 2510 | 2398 | 2920 | 1804 | 1529 |
| RL15 |  |  |  |  |  |  |  |  |  |  |  |  |  |  | 2417 | 2348 | 2461 | 2679 | 2115 | 1753 |
| RL16 |  |  |  |  |  |  |  |  |  |  |  |  |  |  |  | 2212 | 2532 | 2474 | 2116 | 1925 |
| RL17 |  |  |  |  |  |  |  |  |  |  |  |  |  |  |  |  | 2127 | 2918 | 2097 | 1735 |
| RL18 |  |  |  |  |  |  |  |  |  |  |  |  |  |  |  |  |  | 2484 | 2043 | 1867 |
| RL19 |  |  |  |  |  |  |  |  |  |  |  |  |  |  |  |  |  |  | 2096 | 1767 |
| RL20 |  |  |  |  |  |  |  |  |  |  |  |  |  |  |  |  |  |  |  | 1683 |
| RL21 |  |  |  |  |  |  |  |  |  |  |  |  |  |  |  |  |  |  |  |  |
|  | 2520 | 2740 | 2370 | 2439 | 2191 | 2612 | 2726 | 2348 | 2424 | 2633 | 2463 | 2349 | 2453 | 2422 | 2306 | 2447 | 2328 | 2689 | 2036 | 1738 |

**Table 4:** Matched Alignment Sizes for Small Ribosomal Subunit, at threshold 0 (matching by uniqueness)

|  | RL2 | RL3 | RL4 | RL5 | RL6 | RL7 | RL8 | RL9 | RL10 | RL11 | RL12 | RL13 | RL14 | RL15 | RL16 | RL17 | RL18 | RL19 | RL20 | RL21 |
| --- | --- | --- | --- | --- | --- | --- | --- | --- | --- | --- | --- | --- | --- | --- | --- | --- | --- | --- | --- | --- |
| RL2 |  | 2594 | 2143 | 2343 | 2149 | 2608 | 2611 | 2342 | 2333 | 2592 | 2379 | 2095 | 2256 | 2533 | 2318 | 2303 | 2311 | 2599 | 2051 | 1692 |
| RL3 |  |  | 2219 | 2373 | 2371 | 2615 | 2628 | 2363 | 2348 | 2579 | 2406 | 2097 | 2267 | 2535 | 2506 | 2341 | 2444 | 2656 | 2057 | 1871 |
| RL4 |  |  |  | 1895 | 1722 | 2178 | 2140 | 1893 | 1888 | 2117 | 2010 | 1707 | 1886 | 2072 | 1877 | 1858 | 1877 | 2146 | 1653 | 1394 |
| RL5 |  |  |  |  | 2156 | 2356 | 2364 | 2344 | 2333 | 2322 | 2156 | 2078 | 1984 | 2313 | 2160 | 2320 | 2084 | 2319 | 2069 | 1707 |
| RL6 |  |  |  |  |  | 2135 | 2189 | 2153 | 2146 | 2134 | 1960 | 2063 | 1840 | 2138 | 2376 | 2150 | 2251 | 2180 | 2071 | 1879 |
| RL7 |  |  |  |  |  |  | 2617 | 2327 | 2326 | 2596 | 2494 | 2088 | 2267 | 2536 | 2304 | 2304 | 2310 | 2605 | 2043 | 1665 |
| RL8 |  |  |  |  |  |  |  | 2338 | 2341 | 2623 | 2379 | 2113 | 2302 | 2570 | 2385 | 2336 | 2333 | 2669 | 2057 | 1743 |
| RL9 |  |  |  |  |  |  |  |  | 2323 | 2324 | 2156 | 2071 | 1996 | 2315 | 2155 | 2303 | 2102 | 2320 | 2057 | 1700 |
| RL10 |  |  |  |  |  |  |  |  |  | 2327 | 2153 | 2090 | 1996 | 2301 | 2159 | 2302 | 2096 | 2330 | 2055 | 1693 |
| RL11 |  |  |  |  |  |  |  |  |  |  | 2386 | 2091 | 2280 | 2559 | 2318 | 2291 | 2318 | 2596 | 2040 | 1685 |
| RL12 |  |  |  |  |  |  |  |  |  |  |  | 1920 | 2145 | 2324 | 2094 | 2120 | 2069 | 2395 | 1866 | 1508 |
| RL13 |  |  |  |  |  |  |  |  |  |  |  |  | 1806 | 2077 | 2091 | 2052 | 2054 | 2086 | 2003 | 1661 |
| RL14 |  |  |  |  |  |  |  |  |  |  |  |  |  | 2213 | 2037 | 1980 | 2109 | 2290 | 1735 | 1485 |
| RL15 |  |  |  |  |  |  |  |  |  |  |  |  |  |  | 2316 | 2287 | 2304 | 2539 | 2043 | 1697 |
| RL16 |  |  |  |  |  |  |  |  |  |  |  |  |  |  |  | 2149 | 2451 | 2373 | 2066 | 1877 |
| RL17 |  |  |  |  |  |  |  |  |  |  |  |  |  |  |  |  | 2077 | 2321 | 2047 | 1687 |
| RL18 |  |  |  |  |  |  |  |  |  |  |  |  |  |  |  |  |  | 2308 | 1998 | 1827 |
| RL19 |  |  |  |  |  |  |  |  |  |  |  |  |  |  |  |  |  |  | 2033 | 1734 |
| RL20 |  |  |  |  |  |  |  |  |  |  |  |  |  |  |  |  |  |  |  | 1617 |
| RL21 |  |  |  |  |  |  |  |  |  |  |  |  |  |  |  |  |  |  |  |  |
|  | 2329 | 2383 | 1930 | 2193 | 2109 | 2335 | 2355 | 2189 | 2186 | 2325 | 2154 | 2013 | 2046 | 2299 | 2211 | 2170 | 2175 | 2342 | 1977 | 1691 |

**Table 5:** Matched Alignment Sizes for Large Ribosomal Subunit, at threshold 100

|  | RL2 | RL3 | RL4 | RL5 | RL6 | RL9 | RL11 | RL13 | RL14 | RL15 | RL16 | RL17 | RL18 | RL19 | RL20 | RL21 | RL22 | RL23 | RL24 | RL25 | RL27 | RL28 | RL29 | RL30 | RL32 | RL33 | RL34 | RL35 | RL36 |
| --- | --- | --- | --- | --- | --- | --- | --- | --- | --- | --- | --- | --- | --- | --- | --- | --- | --- | --- | --- | --- | --- | --- | --- | --- | --- | --- | --- | --- | --- |
| RL2 |  | 2699 | 2720 | 2875 | 2824 | 2142 | 2505 | 2461 | 3077 | 2658 | 3101 | 2438 | 2672 | 2190 | 2509 | 2174 | 2957 | 3075 | 2435 | 1739 | 2164 | 1932 | 2904 | 2471 | 2296 | 2163 | 1970 | 2094 | 2328 |
| RL3 |  |  | 2789 | 2626 | 2923 | 2149 | 2382 | 2395 | 2873 | 2604 | 2626 | 2456 | 2649 | 2184 | 2161 | 2164 | 3132 | 2661 | 2338 | 1733 | 2184 | 1964 | 2591 | 2290 | 2033 | 1902 | 1993 | 2108 | 1984 |
| RL4 |  |  |  | 2639 | 2709 | 2167 | 2407 | 2418 | 2637 | 2676 | 2647 | 2438 | 2871 | 2209 | 2167 | 2168 | 2788 | 2695 | 2805 | 1747 | 2195 | 1962 | 2652 | 2333 | 2040 | 1894 | 2001 | 2134 | 2011 |
| RL5 |  |  |  |  | 2902 | 2232 | 2492 | 2498 | 2799 | 2692 | 3134 | 2608 | 2775 | 2312 | 2327 | 2309 | 2688 | 3035 | 2483 | 1773 | 2299 | 2014 | 2744 | 2389 | 2164 | 1945 | 2084 | 2203 | 2136 |
| RL6 |  |  |  |  |  | 2216 | 2551 | 2506 | 3043 | 2768 | 2839 | 2651 | 2828 | 2283 | 2277 | 2275 | 2990 | 2773 | 2495 | 1785 | 2286 | 2005 | 2828 | 2455 | 2114 | 1937 | 2039 | 2207 | 2101 |
| RL9 |  |  |  |  |  |  | 2154 | 2156 | 2168 | 2161 | 2174 | 2191 | 2238 | 2283 | 2224 | 2237 | 2153 | 2165 | 502 | 1792 | 2259 | 2025 | 2230 | 1877 | 2099 | 1810 | 2106 | 2190 | 1768 |
| RL11 |  |  |  |  |  |  |  | 2422 | 2492 | 2375 | 2468 | 2223 | 2499 | 2217 | 2179 | 2174 | 2370 | 2539 | 2314 | 1732 | 2187 | 1973 | 2457 | 2131 | 2040 | 2024 | 1991 | 2133 | 1777 |
| RL13 |  |  |  |  |  |  |  |  | 2491 | 2482 | 2498 | 2246 | 2493 | 2208 | 2197 | 2198 | 2340 | 2482 | 1127 | 1755 | 2217 | 1980 | 2445 | 2110 | 2053 | 1852 | 1999 | 2155 | 1800 |
| RL14 |  |  |  |  |  |  |  |  |  | 2643 | 3080 | 2465 | 2752 | 2232 | 2574 | 2227 | 3166 | 3012 | 2328 | 1735 | 2208 | 1989 | 2606 | 2241 | 2345 | 2181 | 2003 | 2126 | 2345 |
| RL15 |  |  |  |  |  |  |  |  |  |  | 2616 | 2509 | 2740 | 2189 | 2169 | 2160 | 2714 | 2700 | 2354 | 1760 | 2196 | 1964 | 2706 | 2388 | 2024 | 1848 | 1970 | 2109 | 2040 |
| RL16 |  |  |  |  |  |  |  |  |  |  |  | 2488 | 2730 | 2240 | 2564 | 2229 | 2812 | 3348 | 2314 | 1759 | 2213 | 1991 | 2610 | 2259 | 2372 | 2191 | 2012 | 2142 | 2325 |
| RL17 |  |  |  |  |  |  |  |  |  |  |  |  | 2755 | 2385 | 2176 | 2341 | 2465 | 2530 | 2207 | 1726 | 2380 | 2146 | 2689 | 2180 | 2190 | 1917 | 2131 | 2181 | 2302 |
| RL18 |  |  |  |  |  |  |  |  |  |  |  |  |  | 2422 | 2223 | 2369 | 2734 | 2739 | 2934 | 1772 | 2417 | 2170 | 2886 | 2454 | 2227 | 1975 | 2176 | 2216 | 2193 |
| RL19 |  |  |  |  |  |  |  |  |  |  |  |  |  |  | 2331 | 2437 | 2188 | 2262 | 580 | 1774 | 2507 | 2277 | 2434 | 1913 | 2361 | 1906 | 2225 | 2315 | 1948 |
| RL20 |  |  |  |  |  |  |  |  |  |  |  |  |  |  |  | 2311 | 2483 | 2518 | 411 | 1787 | 2297 | 2011 | 2248 | 1868 | 2450 | 2161 | 2048 | 2477 | 2152 |
| RL21 |  |  |  |  |  |  |  |  |  |  |  |  |  |  |  |  | 2202 | 2242 | 542 | 1754 | 2692 | 2163 | 2380 | 1887 | 2258 | 1890 | 2177 | 2259 | 1913 |
| RL22 |  |  |  |  |  |  |  |  |  |  |  |  |  |  |  |  |  | 2942 | 2380 | 1739 | 2208 | 1970 | 2595 | 2251 | 2297 | 2160 | 1989 | 2120 | 2294 |
| RL23 |  |  |  |  |  |  |  |  |  |  |  |  |  |  |  |  |  |  | 2405 | 1748 | 2254 | 2007 | 2727 | 2397 | 2381 | 2221 | 2044 | 2152 | 2337 |
| RL24 |  |  |  |  |  |  |  |  |  |  |  |  |  |  |  |  |  |  |  | 391 | 503 | 528 | 2459 | 2093 | 449 | 1111 | 522 | 437 | 1468 |
| RL25 |  |  |  |  |  |  |  |  |  |  |  |  |  |  |  |  |  |  |  |  | 1770 | 1595 | 1745 | 1547 | 1649 | 1564 | 1598 | 1761 | 1362 |
| RL27 |  |  |  |  |  |  |  |  |  |  |  |  |  |  |  |  |  |  |  |  |  | 2234 | 2427 | 1915 | 2300 | 1928 | 2232 | 2295 | 1931 |
| RL28 |  |  |  |  |  |  |  |  |  |  |  |  |  |  |  |  |  |  |  |  |  |  | 2148 | 1719 | 2185 | 1935 | 2015 | 2039 | 1750 |
| RL29 |  |  |  |  |  |  |  |  |  |  |  |  |  |  |  |  |  |  |  |  |  |  |  | 2584 | 2223 | 1957 | 2163 | 2251 | 2160 |
| RL30 |  |  |  |  |  |  |  |  |  |  |  |  |  |  |  |  |  |  |  |  |  |  |  |  | 1765 | 1579 | 1732 | 1851 | 1738 |
| RL32 |  |  |  |  |  |  |  |  |  |  |  |  |  |  |  |  |  |  |  |  |  |  |  |  |  | 2183 | 2074 | 2130 | 2132 |
| RL33 |  |  |  |  |  |  |  |  |  |  |  |  |  |  |  |  |  |  |  |  |  |  |  |  |  |  | 1741 | 1819 | 1921 |
| RL34 |  |  |  |  |  |  |  |  |  |  |  |  |  |  |  |  |  |  |  |  |  |  |  |  |  |  |  | 2089 | 1779 |
| RL35 |  |  |  |  |  |  |  |  |  |  |  |  |  |  |  |  |  |  |  |  |  |  |  |  |  |  |  |  | 1800 |
| RL36 |  |  |  |  |  |  |  |  |  |  |  |  |  |  |  |  |  |  |  |  |  |  |  |  |  |  |  |  |  |
|  | 2485 | 2378 | 2390 | 2471 | 2486 | 2067 | 2257 | 2214 | 2494 | 2365 | 2492 | 2336 | 2497 | 2172 | 2189 | 2148 | 2469 | 2514 | 1604 | 1664 | 2168 | 1953 | 2459 | 2086 | 2101 | 1918 | 1961 | 2064 | 1993 |

**Table 6:** Matched Alignment Sizes for Large Ribosomal Subunit, at threshold 0 (matching by uniqueness)

|  | RL2 | RL3 | RL4 | RL5 | RL6 | RL9 | RL11 | RL13 | RL14 | RL15 | RL16 | RL17 | RL18 | RL19 | RL20 | RL21 | RL22 | RL23 | RL24 | RL25 | RL27 | RL28 | RL29 | RL30 | RL32 | RL33 | RL34 | RL35 | RL36 |
| --- | --- | --- | --- | --- | --- | --- | --- | --- | --- | --- | --- | --- | --- | --- | --- | --- | --- | --- | --- | --- | --- | --- | --- | --- | --- | --- | --- | --- | --- |
| RL2 |  | 2144 | 2173 | 2333 | 2307 | 2079 | 2277 | 2286 | 2568 | 2115 | 2547 | 2041 | 2313 | 2120 | 2325 | 2095 | 2325 | 2552 | 217 | 1698 | 2100 | 1895 | 2257 | 1964 | 2182 | 1875 | 1918 | 2052 | 1900 |
| RL3 |  |  | 2139 | 2179 | 2162 | 2087 | 2152 | 2169 | 2165 | 2102 | 2168 | 2033 | 2178 | 2126 | 2109 | 2095 | 2112 | 2177 | 177 | 1703 | 2115 | 1899 | 2140 | 1851 | 1965 | 1651 | 1933 | 2068 | 1648 |
| RL4 |  |  |  | 2205 | 2188 | 2099 | 2175 | 2191 | 2190 | 2118 | 2197 | 2045 | 2207 | 2140 | 2119 | 2102 | 2117 | 2193 | 189 | 1704 | 2126 | 1912 | 2170 | 1865 | 1978 | 1663 | 1944 | 2075 | 1652 |
| RL5 |  |  |  |  | 2425 | 2176 | 2316 | 2319 | 2388 | 2151 | 2370 | 2162 | 2379 | 2255 | 2257 | 2221 | 2150 | 2369 | 221 | 1735 | 2235 | 1960 | 2394 | 2038 | 2093 | 1727 | 2005 | 2161 | 1771 |
| RL6 |  |  |  |  |  | 2164 | 2307 | 2310 | 2344 | 2149 | 2337 | 2134 | 2368 | 2221 | 2214 | 2187 | 2130 | 2324 | 221 | 1725 | 2204 | 1949 | 2379 | 2016 | 2045 | 1694 | 1981 | 2144 | 1713 |
| RL9 |  |  |  |  |  |  | 2088 | 2106 | 2110 | 2095 | 2122 | 2142 | 2178 | 2232 | 2176 | 2187 | 2102 | 2119 | 167 | 1735 | 2224 | 1967 | 2181 | 1824 | 2059 | 1697 | 2018 | 2147 | 1720 |
| RL11 |  |  |  |  |  |  |  | 2305 | 2317 | 2117 | 2300 | 2061 | 2319 | 2143 | 2129 | 2115 | 2109 | 2299 | 219 | 1693 | 2133 | 1922 | 2289 | 1978 | 1994 | 1672 | 1938 | 2074 | 1668 |
| RL13 |  |  |  |  |  |  |  |  | 2312 | 2130 | 2306 | 2064 | 2323 | 2152 | 2141 | 2129 | 2121 | 2305 | 205 | 1713 | 2145 | 1918 | 2292 | 1984 | 1988 | 1670 | 1952 | 2090 | 1668 |
| RL14 |  |  |  |  |  |  |  |  |  | 2146 | 2600 | 2089 | 2349 | 2165 | 2392 | 2155 | 2388 | 2606 | 217 | 1710 | 2151 | 1940 | 2318 | 2012 | 2259 | 1941 | 1961 | 2085 | 1998 |
| RL15 |  |  |  |  |  |  |  |  |  |  | 2166 | 2062 | 2162 | 2137 | 2120 | 2107 | 2120 | 2132 | 181 | 1713 | 2127 | 1917 | 2110 | 1842 | 1975 | 1657 | 1936 | 2073 | 1653 |
| RL16 |  |  |  |  |  |  |  |  |  |  |  | 2089 | 2335 | 2171 | 2370 | 2144 | 2347 | 2539 | 216 | 1724 | 2155 | 1935 | 2304 | 2001 | 2226 | 1902 | 1963 | 2095 | 1931 |
| RL17 |  |  |  |  |  |  |  |  |  |  |  |  | 2302 | 2346 | 2146 | 2280 | 2049 | 2121 | 222 | 1677 | 2337 | 2099 | 2305 | 1798 | 2155 | 1724 | 2094 | 2144 | 1802 |
| RL18 |  |  |  |  |  |  |  |  |  |  |  |  |  | 2366 | 2177 | 2310 | 2144 | 2381 | 293 | 1731 | 2366 | 2127 | 2521 | 2056 | 2177 | 1774 | 2126 | 2166 | 1821 |
| RL19 |  |  |  |  |  |  |  |  |  |  |  |  |  |  | 2260 | 2370 | 2138 | 2208 | 235 | 1748 | 2438 | 2156 | 2392 | 1875 | 2248 | 1816 | 2190 | 2260 | 1889 |
| RL20 |  |  |  |  |  |  |  |  |  |  |  |  |  |  |  | 2231 | 2345 | 2379 | 174 | 1737 | 2226 | 1960 | 2197 | 1840 | 2294 | 1952 | 2003 | 2177 | 2011 |
| RL21 |  |  |  |  |  |  |  |  |  |  |  |  |  |  |  |  | 2125 | 2179 | 224 | 1720 | 2367 | 2089 | 2317 | 1838 | 2191 | 1777 | 2130 | 2203 | 1849 |
| RL22 |  |  |  |  |  |  |  |  |  |  |  |  |  |  |  |  |  | 2373 | 170 | 1707 | 2132 | 1917 | 2107 | 1810 | 2215 | 1908 | 1946 | 2081 | 1936 |
| RL23 |  |  |  |  |  |  |  |  |  |  |  |  |  |  |  |  |  |  | 227 | 1716 | 2187 | 1955 | 2351 | 2006 | 2289 | 1957 | 1988 | 2107 | 1999 |
| RL24 |  |  |  |  |  |  |  |  |  |  |  |  |  |  |  |  |  |  |  | 116 | 238 | 211 | 288 | 169 | 224 | 180 | 207 | 182 | 195 |
| RL25 |  |  |  |  |  |  |  |  |  |  |  |  |  |  |  |  |  |  |  |  | 1733 | 1539 | 1713 | 1517 | 1602 | 1474 | 1566 | 1704 | 1323 |
| RL27 |  |  |  |  |  |  |  |  |  |  |  |  |  |  |  |  |  |  |  |  |  | 2164 | 2376 | 1863 | 2243 | 1814 | 2194 | 2236 | 1879 |
| RL28 |  |  |  |  |  |  |  |  |  |  |  |  |  |  |  |  |  |  |  |  |  |  | 2109 | 1654 | 2036 | 1697 | 1980 | 1989 | 1685 |
| RL29 |  |  |  |  |  |  |  |  |  |  |  |  |  |  |  |  |  |  |  |  |  |  |  | 2052 | 2188 | 1771 | 2132 | 2210 | 1835 |
| RL30 |  |  |  |  |  |  |  |  |  |  |  |  |  |  |  |  |  |  |  |  |  |  |  |  | 1724 | 1427 | 1711 | 1821 | 1441 |
| RL32 |  |  |  |  |  |  |  |  |  |  |  |  |  |  |  |  |  |  |  |  |  |  |  |  |  | 1988 | 2048 | 2084 | 2033 |
| RL33 |  |  |  |  |  |  |  |  |  |  |  |  |  |  |  |  |  |  |  |  |  |  |  |  |  |  | 1655 | 1708 | 1693 |
| RL34 |  |  |  |  |  |  |  |  |  |  |  |  |  |  |  |  |  |  |  |  |  |  |  |  |  |  |  | 2051 | 1730 |
| RL35 |  |  |  |  |  |  |  |  |  |  |  |  |  |  |  |  |  |  |  |  |  |  |  |  |  |  |  |  | 1763 |
| RL36 |  |  |  |  |  |  |  |  |  |  |  |  |  |  |  |  |  |  |  |  |  |  |  |  |  |  |  |  |  |
|  | 2095 | 1980 | 1996 | 2107 | 2084 | 2000 | 2040 | 2046 | 2138 | 1975 | 2127 | 2019 | 2141 | 2100 | 2088 | 2062 | 2040 | 2144 | 207 | 1613 | 2090 | 1878 | 2132 | 1785 | 2018 | 1695 | 1904 | 1998 | 1722 |

In order to produce an interaction score for the two proteins, we run the PLM algorithm [12] on the concatenated alignments. This results in a list of residue pairs of the alignment ordered by their interaction strength. We filtered out the pairs that contain one residue of one protein and one of the other. This results in a list of *possibly* interacting inter-protein residue pairs ordered by the interaction score. In order to arrive at an interaction score for the two proteins we took the mean of the scores for the 4 highest scoring pairs (PPI-score). The number 4 was used because it performed best on the small ribosomal subunit, but the predictive performance on a larger-scale network is virtually identical for any value between 1 and 6 (see Figure S8). The list of protein pairs ordered by this score was used for prediction. The first few predictions are shown in Table 8. For completeness, we show the same table but with the score calculated by the Gaussian approximation of [2] in Table 9. Finally in Table 11 we display for the LSU the number of intra/inter-protein contacts, while in Table 12 we do the same for the LRU.

Table 10 shows the interaction scores for the protein pairs of the Trp Operon.

### 4.1 Structural view of the Ribosomal Complex

In Fig. 10 we display a cartoon view of the ribosomal protein network. The contact map for the the small and large ribosomal units are displayed in Fig. 11

**Table 7:** Matched Alignment Sizes for Trp for different matching thresholds (threshold 0 corresponds to matching by uniqueness)

| P1 | P2 | tr=100 | tr=0 |
| --- | --- | --- | --- |
| TrpC | TrpG | 4272 | 18 |
| TrpE | TrpF | 2519 | 830 |
| TrpA | TrpD | 2823 | 743 |
| TrpD | TrpG | 6249 | 28 |
| TrpB | TrpF | 3643 | 95 |
| TrpB | TrpD | 3737 | 95 |
| TrpB | TrpG | 8053 | 41 |
| TrpE | TrpG | 5324 | 8 |
| TrpD | TrpF | 2819 | 695 |
| TrpC | TrpF | 3825 | 1578 |
| TrpA | TrpC | 3198 | 1546 |
| TrpC | TrpD | 3392 | 748 |
| TrpA | TrpF | 3357 | 1433 |
| TrpA | TrpE | 3118 | 905 |
| TrpD | TrpE | 2681 | 482 |
| TrpB | TrpC | 3326 | 82 |
| TrpB | TrpE | 3911 | 53 |
| TrpC | TrpE | 2976 | 930 |
| TrpF | TrpG | 3635 | 32 |
| TrpA | TrpB | 4374 | 95 |
| TrpA | TrpG | 4646 | 22 |

**Figure 9:**
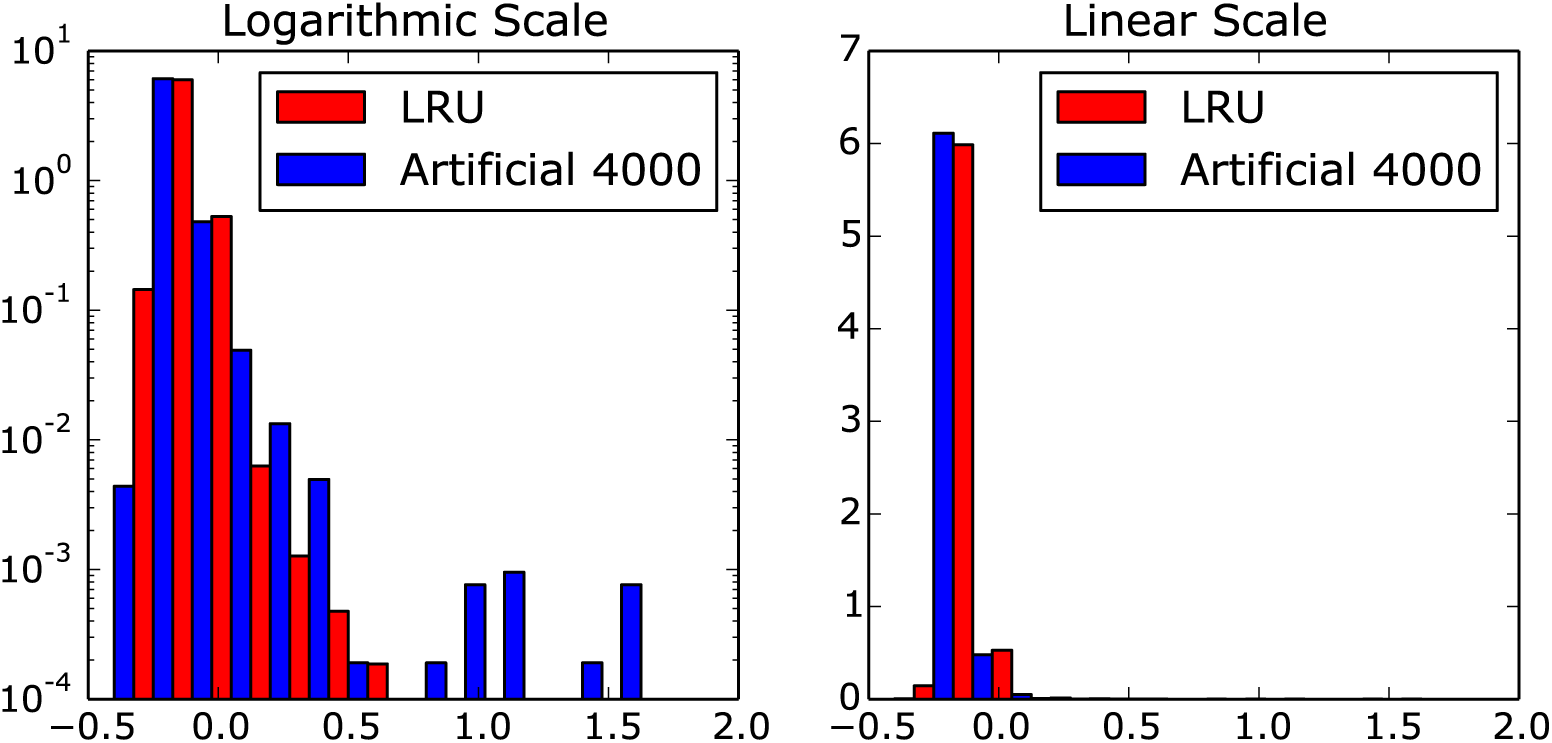
Histograms of interaction scores resulting from the analysis of the LRU and the artificial complex (combined strategy). Both intra- and inter-protein scores are included. The plots are normalized such that the area of all bars of a given color sums to one. The data is shown both on a logarithmic (left) and on a linear scale (right).

**Table 8:** Ordered List of Interaction Candidates SRU (left) and LRU (right) based on plmDCA scores; the fourth column indicates whether the protein pair is indeed interacting

| P1 | P2 | Score | Interacting | P1 | P2 | Score | Interacting |
| --- | --- | --- | --- | --- | --- | --- | --- |
| RS10 | RS14 | 0.618890 | 1 | RL20 | RL21 | 0.576795 | 1 |
| RS18 | RS6 | 0.422457 | 1 | RL14 | RL19 | 0.514107 | 1 |
| RS14 | RS3 | 0.394753 | 1 | RL15 | RL35 | 0.440323 | 1 |
| RS10 | RS9 | 0.347508 | 1 | RL15 | RL21 | 0.439233 | 1 |
| RS13 | RS19 | 0.317640 | 1 | RL17 | RL32 | 0.425920 | 1 |
| RS13 | RS21 | 0.306248 | 0 | RL20 | RL32 | 0.421733 | 1 |
| RS11 | RS21 | 0.296700 | 1 | RL23 | RL29 | 0.414060 | 1 |
| RS14 | RS19 | 0.291335 | 1 | RL13 | RL20 | 0.334348 | 1 |
| RS12 | RS21 | 0.290965 | 0 | RL19 | RL3 | 0.328640 | 1 |
| RS16 | RS4 | 0.287438 | 0 | RL30 | RL34 | 0.326368 | 0 |
| RS21 | RS7 | 0.287102 | 0 | RL22 | RL32 | 0.324540 | 1 |
| RS13 | RS15 | 0.284783 | 0 | RL16 | RL36 | 0.318915 | 1 |
| RS12 | RS16 | 0.283105 | 0 | RL16 | RL33 | 0.313083 | 0 |
| RS19 | RS21 | 0.282142 | 0 | RL33 | RL36 | 0.307188 | 0 |
| RS10 | RS18 | 0.279595 | 0 | RL27 | RL34 | 0.306283 | 0 |

**Table 9:** Ordered List of Interaction Candidates SRU (left) and LRU (right) based on Gaussian scores; the fourth column indicates whether the protein pair is indeed interacting

| P1 | P2 | Score | Interacting | P1 | P2 | Score | Interacting |
| --- | --- | --- | --- | --- | --- | --- | --- |
| RS10 | RS9 | 1.123465 | 1 | RL20 | RL21 | 1.665182 | 1 |
| RS10 | RS14 | 1.102428 | 1 | RL14 | RL19 | 1.430611 | 1 |
| RS12 | RS21 | 1.079407 | 0 | RL15 | RL21 | 1.333611 | 1 |
| RS13 | RS18 | 1.029537 | 0 | RL15 | RL35 | 1.134808 | 1 |
| RS14 | RS17 | 1.001716 | 0 | RL23 | RL29 | 1.086992 | 1 |
| RS12 | RS15 | 0.997813 | 0 | RL20 | RL32 | 1.037364 | 1 |
| RS18 | RS6 | 0.963688 | 1 | RL22 | RL32 | 1.029724 | 1 |
| RS11 | RS13 | 0.943144 | 0 | RL30 | RL34 | 1.008776 | 0 |
| RS19 | RS21 | 0.942921 | 0 | RL17 | RL32 | 1.002790 | 1 |
| RS15 | RS18 | 0.938286 | 0 | RL34 | RL36 | 0.983223 | 0 |
| RS14 | RS15 | 0.933949 | 0 | RL21 | RL2 | 0.977507 | 0 |
| RS13 | RS15 | 0.933337 | 0 | RL21 | RL34 | 0.958441 | 0 |
| RS13 | RS19 | 0.918528 | 1 | RL18 | RL34 | 0.942494 | 0 |
| RS18 | RS21 | 0.918101 | 1 | RL36 | RL6 | 0.925895 | 1 |
| RS10 | RS13 | 0.917482 | 0 | RL33 | RL36 | 0.898444 | 0 |

**Table 10:** Ordered List of Interaction Scores for the Trp Operon based on plmDCA scores

|  |  |  |
| --- | --- | --- |
| TrpA | TrpB | 0.375 |
| TrpE | TrpG | 0.295 |
| TrpA | TrpC | 0.167 |
| TrpA | TrpF | 0.162 |
| TrpC | TrpF | 0.146 |
| TrpA | TrpD | 0.144 |
| TrpC | TrpD | 0.141 |
| TrpB | TrpF | 0.136 |
| TrpC | TrpE | 0.135 |
| TrpD | TrpF | 0.135 |
| TrpB | TrpC | 0.132 |
| TrpA | TrpE | 0.126 |
| TrpC | TrpG | 0.121 |
| TrpB | TrpD | 0.120 |
| TrpE | TrpF | 0.115 |
| TrpD | TrpE | 0.107 |
| TrpF | TrpG | 0.107 |
| TrpA | TrpG | 0.104 |
| TrpD | TrpG | 0.100 |
| TrpB | TrpE | 0.096 |
| TrpB | TrpG | 0.071 |

**Table 11:** Left table: number of intra-protein contacts below 8Å of all residues (SEP=0 column), and considering only those with a distance on the sequence of at least 5 residues (SEP = 5 column) for the SRU. Right table: number of inter-protein contacts below 8Åfor the SRU. Fractions are defined as 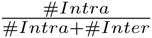 where *#Inter* is computed assuming SEP=0,5 respectively.

| SRU Intra-Protein |  |  |
| --- | --- | --- |
|  | SEP=0 | SEP=5 |
| RS2 | 2337 | 1610 |
| RS3 | 2217 | 1494 |
| RS4 | 1728 | 1152 |
| RS5 | 1684 | 1175 |
| RS6 | 1002 | 666 |
| RS7 | 1494 | 982 |
| RS8 | 1334 | 903 |
| RS9 | 1240 | 799 |
| RS10 | 878 | 557 |
| RS11 | 1220 | 822 |
| RS12 | 1136 | 731 |
| RS13 | 1024 | 623 |
| RS14 | 790 | 440 |
| RS15 | 823 | 489 |
| RS16 | 685 | 436 |
| RS17 | 733 | 487 |
| RS18 | 482 | 293 |
| RS19 | 748 | 482 |
| RS20 | 792 | 464 |
| RS21 | 297 | 110 |
| SUM: | 22644 | 14715 |

Table 11: Left table: number of intra-protein contacts below 8Å of all residues (SEP=0 column), and considering only those with a distance on the sequence of at least 5 residues (SEP = 5 column) for the SRU.
| SRU Inter-Protein |  |  |
| --- | --- | --- |
| RS2 | RS5 | 4 |
| RS2 | RS8 | 3 |
| RS3 | RS5 | 17 |
| RS3 | RS10 | 105 |
| RS3 | RS14 | 209 |
| RS4 | RS5 | 84 |
| RS5 | RS8 | 120 |
| RS6 | RS18 | 150 |
| RS7 | RS9 | 19 |
| RS7 | RS11 | 46 |
| RS8 | RS12 | 12 |
| RS8 | RS17 | 28 |
| RS9 | RS10 | 28 |
| RS9 | RS14 | 7 |
| RS10 | RS14 | 150 |
| RS11 | RS18 | 20 |
| RS11 | RS21 | 199 |
| RS12 | RS17 | 34 |
| RS13 | RS19 | 80 |
| RS14 | RS19 | 50 |
| RS18 | RS21 | 36 |
| SUM: |  | 1401 |
| FRACTION | SEP=0 | 0.058 |
| FRACTION | SEP=5 | 0.087 |

**Table 12:** Left table: number of intra-protein contacts below 8Å of all residues (SEP=0 column), and considering only those with a distance on the sequence of at least 5 residues (SEP = 5 column) for the LRU. Right table: number of inter-protein contacts below 8Åfor the LRU. Fractions are defined as 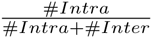 where *#Inter* is computed assuming SEP=0,5 respectively.

| LRU Intra-Protein |  |  | LRU Inter-Protein |  |  |
| --- | --- | --- | --- | --- | --- |
|  | SEP=0 | SEP=5 |  |  |  |
| RL32 | 324 | 157 | RL32 | RL17 | 78 |
| RL33 | 399 | 256 | RL32 | RL20 | 17 |
| RL34 | 303 | 145 | RL32 | RL22 | 73 |
| RL35 | 495 | 268 | RL33 | RL35 | 21 |
| RL36 | 332 | 208 | RL35 | RL15 | 149 |
| RL2 | 2687 | 1801 | RL35 | RL27 | 1 |
| RL3 | 1931 | 1263 | RL36 | RL6 | 10 |
| RL4 | 1869 | 1199 | RL36 | RL16 | 1 |
| RL5 | 1887 | 1257 | RL3 | RL13 | 20 |
| RL6 | 1811 | 1217 | RL3 | RL14 | 34 |
| RL9 | 1360 | 855 | RL3 | RL17 | 21 |
| RL11 | 1390 | 903 | RL3 | RL19 | 123 |
| RL13 | 1464 | 959 | RL4 | RL15 | 83 |
| RL14 | 1266 | 869 | RL4 | RL20 | 6 |
| RL15 | 920 | 481 | RL9 | RL28 | 63 |
| RL16 | 1343 | 915 | RL13 | RL20 | 118 |
| RL17 | 1194 | 767 | RL13 | RL21 | 8 |
| RL18 | 1150 | 777 | RL14 | RL19 | 191 |
| RL19 | 1043 | 669 | RL15 | RL20 | 2 |
| RL20 | 1045 | 600 | RL15 | RL21 | 24 |
| RL21 | 915 | 600 | RL16 | RL25 | 53 |
| RL22 | 1085 | 720 | RL16 | RL27 | 9 |
| RL23 | 735 | 461 | RL17 | RL22 | 12 |
| RL24 | 386 | 233 | RL18 | RL27 | 12 |
| RL25 | 893 | 597 | RL20 | RL21 | 229 |
| RL27 | 692 | 442 | RL23 | RL29 | 81 |
| RL29 | 538 | 303 | SUM: |  | 1439 |
| RL30 | 511 | 321 | FRACTION | SEP=0 | 0.045 |
| RL28 | 587 | 351 | FRACTION | SEP=5 | 0.068 |
| SUM: | 30555 | 19594 |  |  |  |

**Figure 10:**
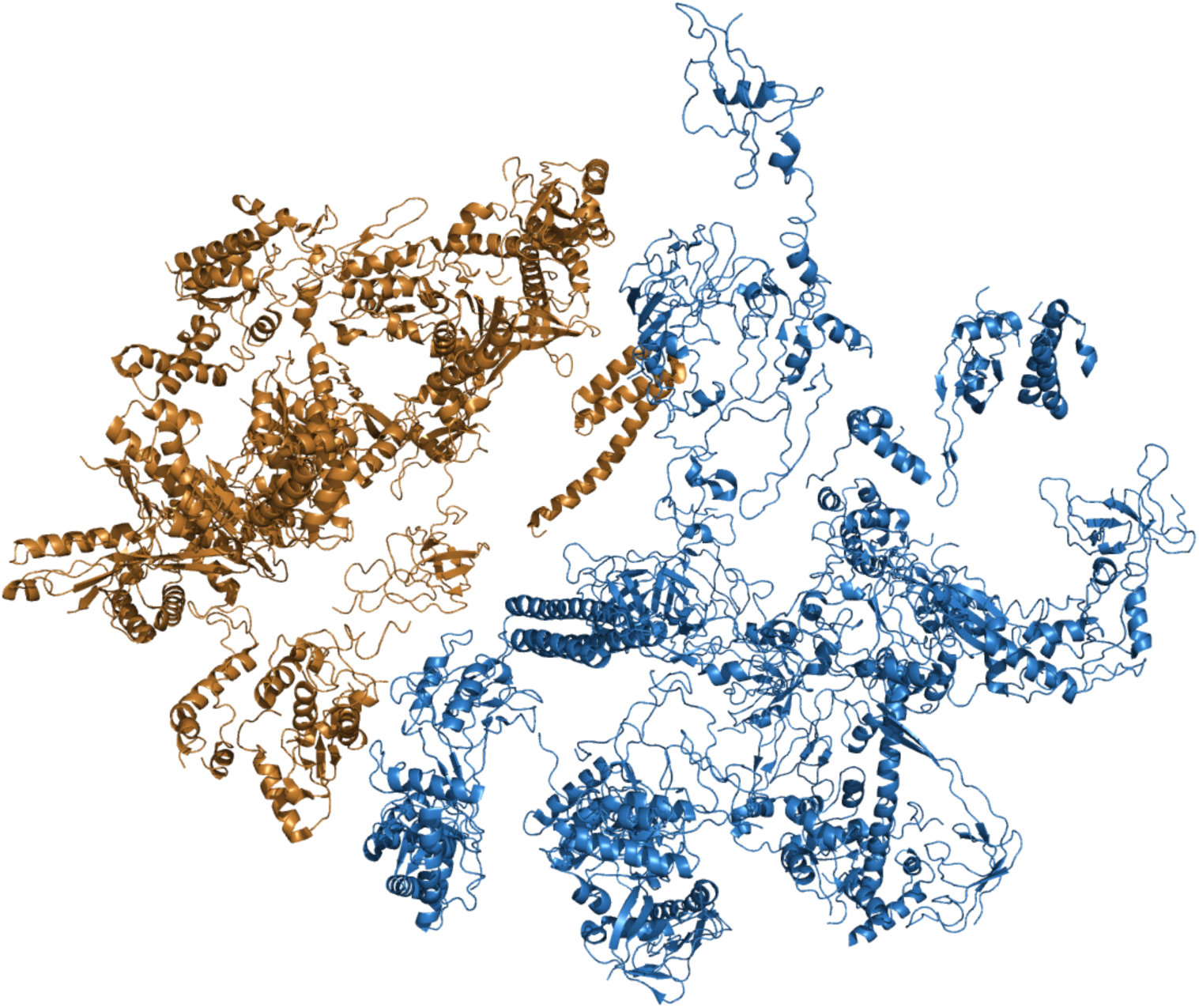
Cartoon view of the small (brass color) and large (blue color) bacterial ribosomal complexes 2Z4K, 2Z4L. For the ease of visualization we have carved out the ribosomal RNAs strands.

**Figure 11:**
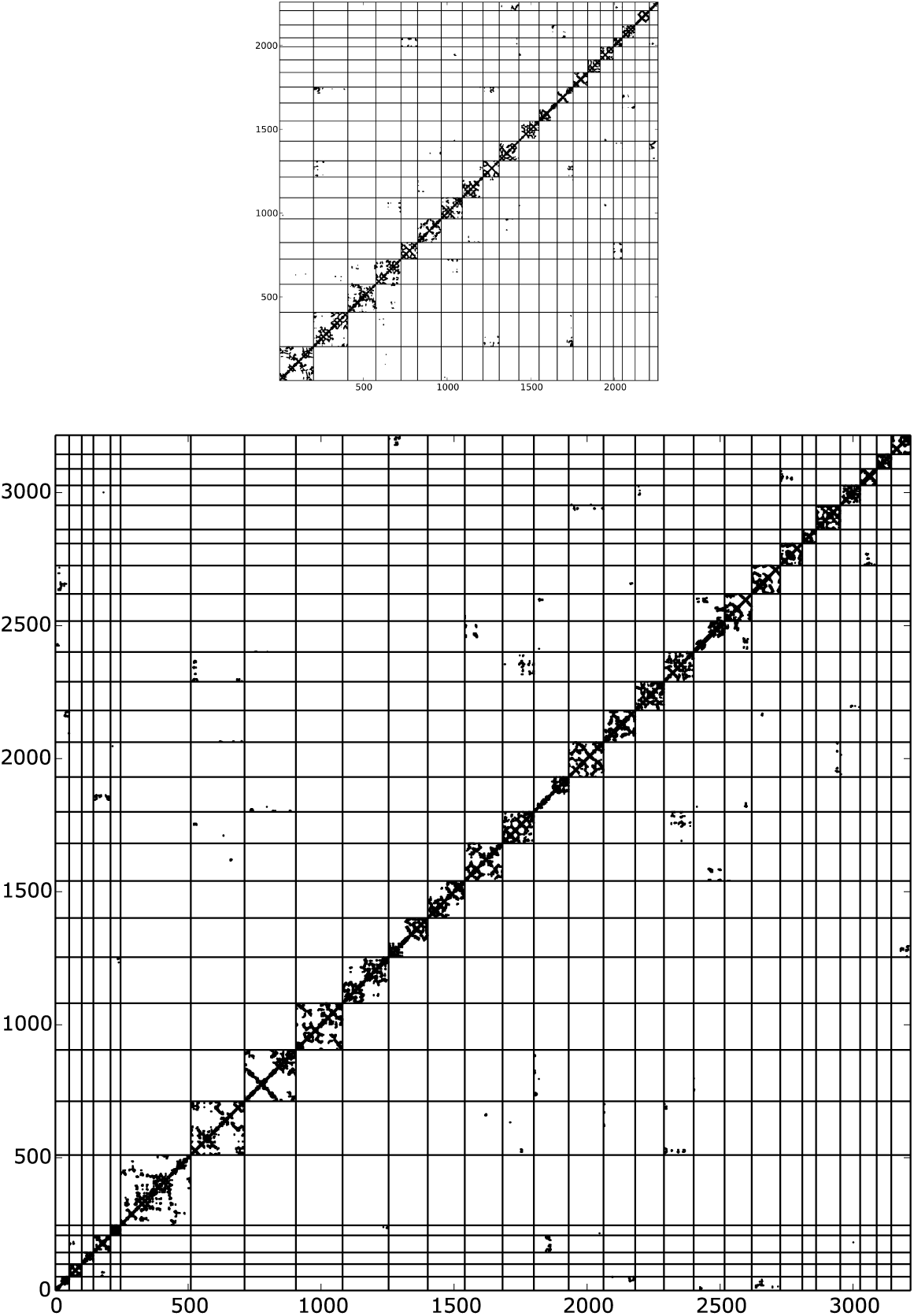
Upper panel: contact map of the SRU (threshold distance 8Å). Lower panel: contact map of the LRU.

## 5 Artificial Data

An artificial large network consisting of 5 proteins was created in two steps:

1) First, a contact map was defined. This contact map contains the information which residues are in contact. This includes internal residue contacts (where both residues belong to one of the 5 proteins) and inter-protein residue contacts (where one residue belongs to one protein and the other to a different protein). The contact map is therefore a binary, symmetric matrix of size *N_all_ × N_all_* with *N_all_* = *N*_1_ + *N*_2_ + *N*_3_ + *N*_4_ + *N*_5_ where *N_i_* is the number of residues in the *i*^th^ protein. We decided to use the Kunitz domain (PF00014) as a model for the proteins and set all *N_i_* = 53. The 53 × 53 submatrices that define the contacts within each protein were defined by extracting the contacts of the PDB structure 5pti of the Kunitz domain. This implies that the internal structure of every protein is the same.

We defined as contacting proteins the protein pairs 1 – 2, 2 – 3, 3 – 4, 4 – 5 and 1 – 5. For the 53 × 53 submatrices that define the contacts between contacting protein pairs we used random binary matrices with 10% of the number of internal contacts. This was done individually for each contacting protein pair such that no two contact matrices between two proteins were the same. For non-contacting protein pairs all entries of the contact matrices were set to 0.

The resulting contact map can be seen in Fig. 12.

**Figure 12:**
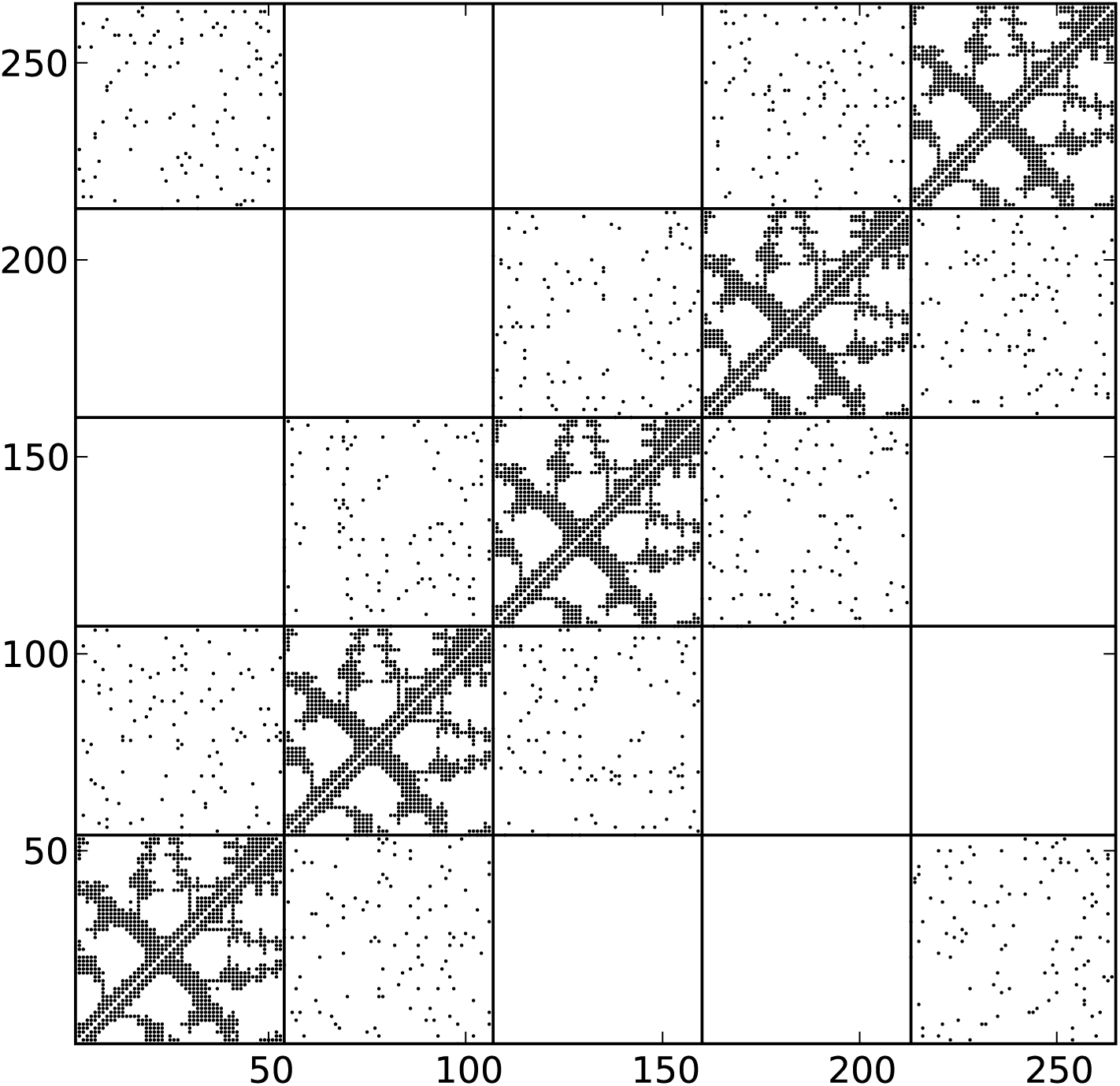
Contact map of the artificial protein complex

2) Couplings for every contact in the contact map were defined. As a basis for this, couplings and fields inferred from the PF00014 PFAM alignment (Kunitz Domain) were used. This inference was done using a masking with the PDB structure, such that only couplings corresponding to PDB-contacts were allowed to differ from zero. Given that the same PDB-contacts were used to define the contacts within one protein in the artificial complex, we could use the couplings thus inferred without change for the couplings within the artificial proteins.

Then we defined the couplings for residue contacts between two proteins. For every such a resiue contact we chose randomly a coupling of an internal contact as inferred from the Kunitz domain alignment and assigned it to the residue contact.

Notice that the ‘coupling’ between two sites *i* and *j* is actually a 21 × 21 matrix *J_ij_* (*a, b*) where *a* and *b* can be any of the 21 amino acids. Given that the internal structure of these matrices might be important we decided to treat the matrices *J_ij_* as single entities and not change their internal structure.

The fields for every residue, a vector of length 21 for every of the 5 · 53 residues, were randomly chosen from the inferred fields.

From these couplings and fields, sequences were generated by MC (see section below) and inferred by plmDCA. Interestingly, a crude comparison between the histogram of the scores in the artificial model seem to be very close to that obtained for instance for the LRU case as shown in Fig. 9.

In Table 13 we compare the ranks of the strongest inter-protein residue interaction scores in the generating model and the inferred model. The first column represents the rank of the inter-protein residue interaction in the generating model, the second column the rank of the same residue interaction in the inferred model. The model was inferred with the combined strategy and with 4000 sequences. The numbering is treating the complex as one large protein.

**Table 13:** Original vs. inferred rank for the 10 largest original inter-protein residue interaction scores and the 10 largest inferred inter-protein residue interaction scores

| Original Rank | Inferred Rank |
| --- | --- |
| 1 | 101 |
| 2 | 13806 |
| 3 | 10658 |
| 4 | 64 |
| 5 | 4 |
| 6 | 9575 |
| 7 | 1 |
| 8 | 15890 |
| 9 | 6712 |
| 10 | 1035 |
| 7 | 1 |
| 32 | 2 |
| 41 | 3 |
| 5 | 4 |
| 11 | 5 |
| 11473 | 6 |
| 22464 | 7 |
| 53 | 8 |
| 1877 | 9 |
| 26 | 10 |

### 5.1 Monte Carlo Sequence Generation

Given the parameters of the artificial model, a simple MCMC algorithm was run to generate samples from the corresponding distribution. We used one million MC steps to equilibrate the chain and took a sample every one million steps.

**Figure 13:**
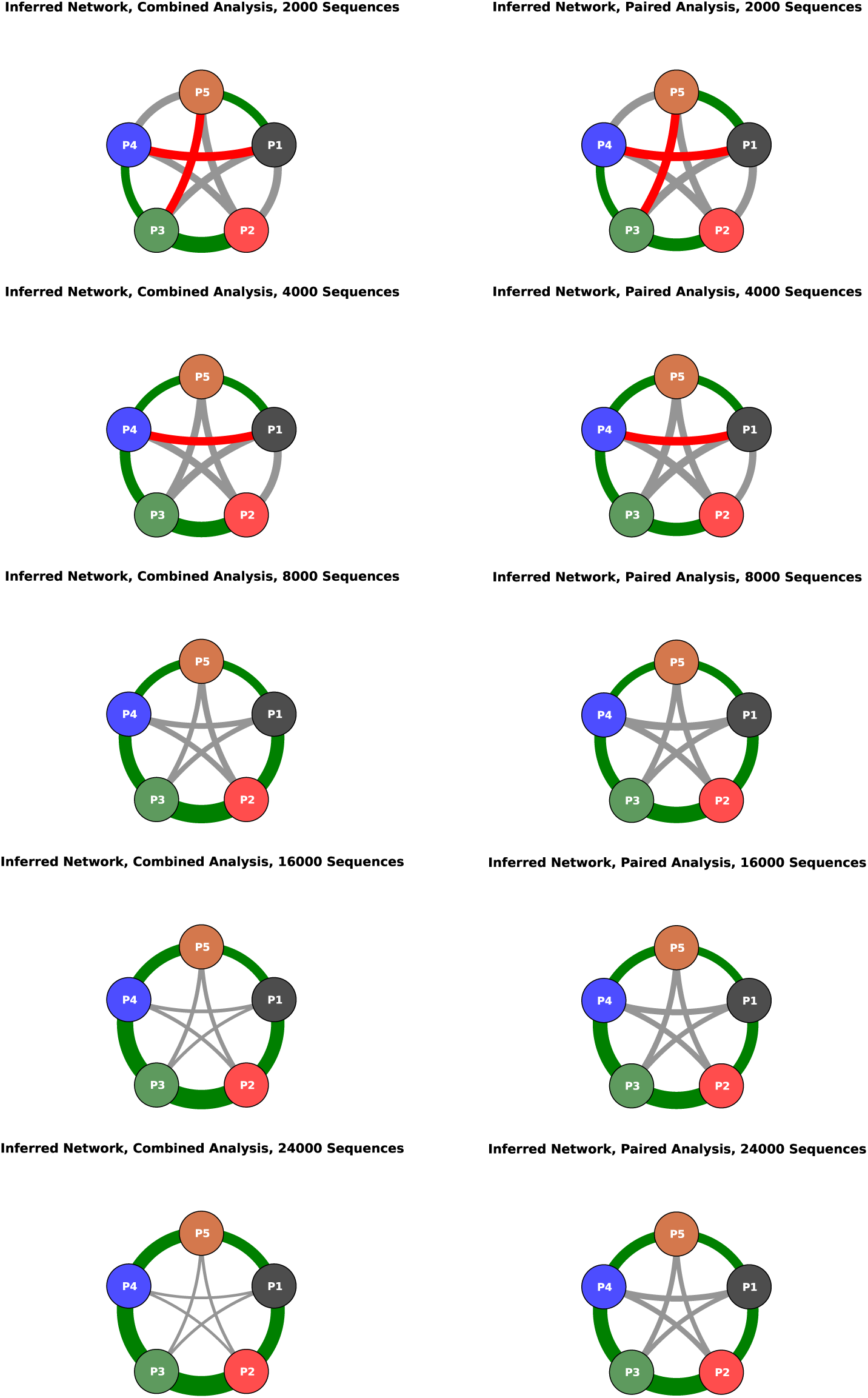
Inferred protein network for different sample sizes; the line-thickness is proportional to the inferred interaction scores between the proteins (mean of the 4 highest residue interaction scores). The thickness has been normalized in the sense that the scores have been divided by the mean of the scores of the network. The color code is applied for the first 5 predictions and shows a green line if the prediction is a true positive and a red line if the prediction is a false positive. Predictions after the first 5 are grey. **Combined Analysis**: The complete sequences in their whole length were used for the inference and calculation of the scores **Paired Anlysis**: Every protein family was independently cut out of the generated sequences and thus a MSA for only this protein created. These single MSAs were then paired for all protein pairs and used for inference and calculation of the scores.

## 6 Large scale network inference

In order to test the approach on a larger scale we created all possible protein pairs from all proteins in the ribosome and the trp operon. The matching procedure was identical to the procedure used in the individual systems.

**Figure 14:**
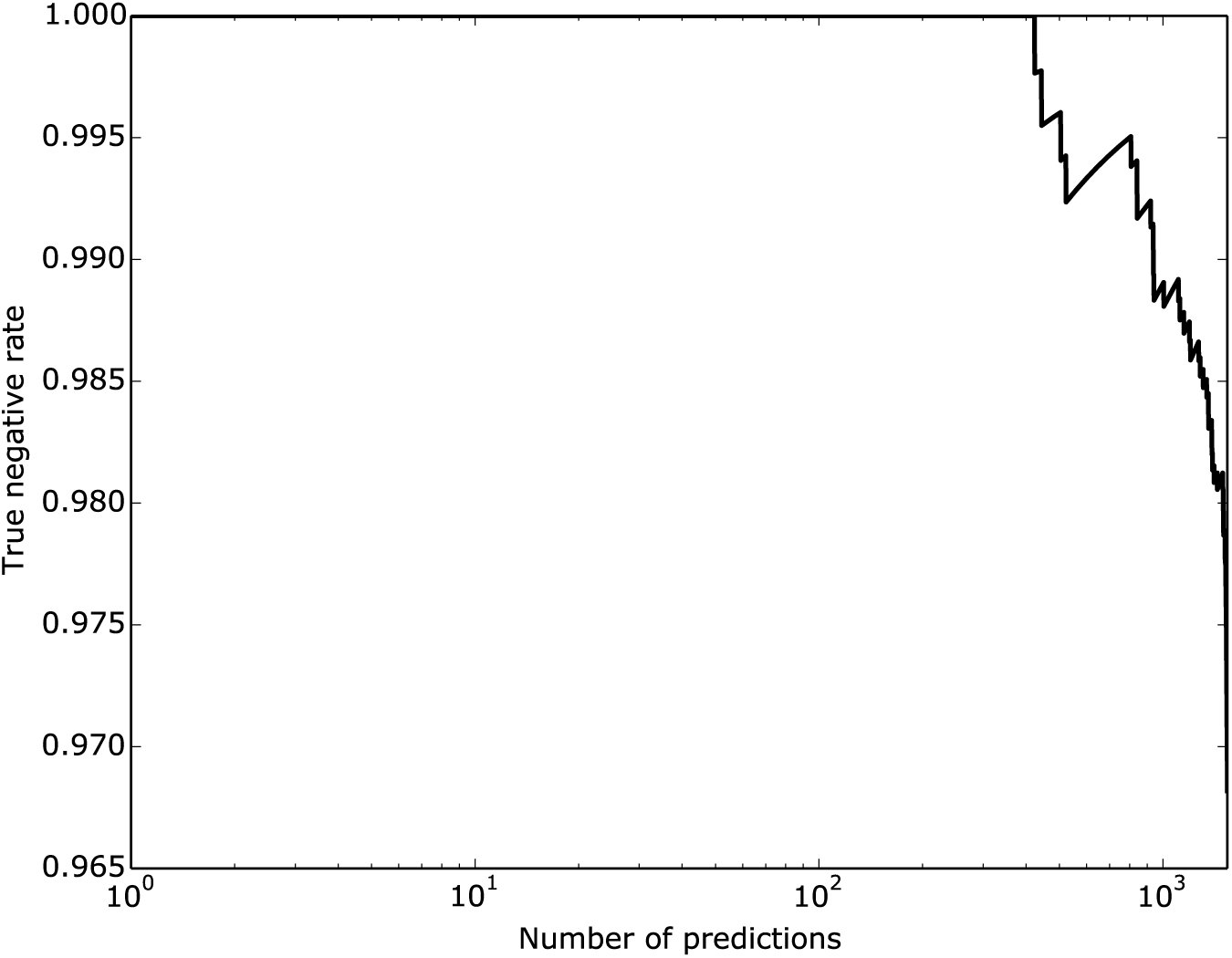
True negative rate; all possible protein pairs between RS,RL and Trp proteins are considered and the protein-protein interaction score is defined as the average of the 4 largest interaction scores on the residue level (as in the main paper). The true negative rate is the fraction of true negatives in the N pairs with the lowest interaction score, where N is the value indicated by the x-axis.

**Figure 15:**
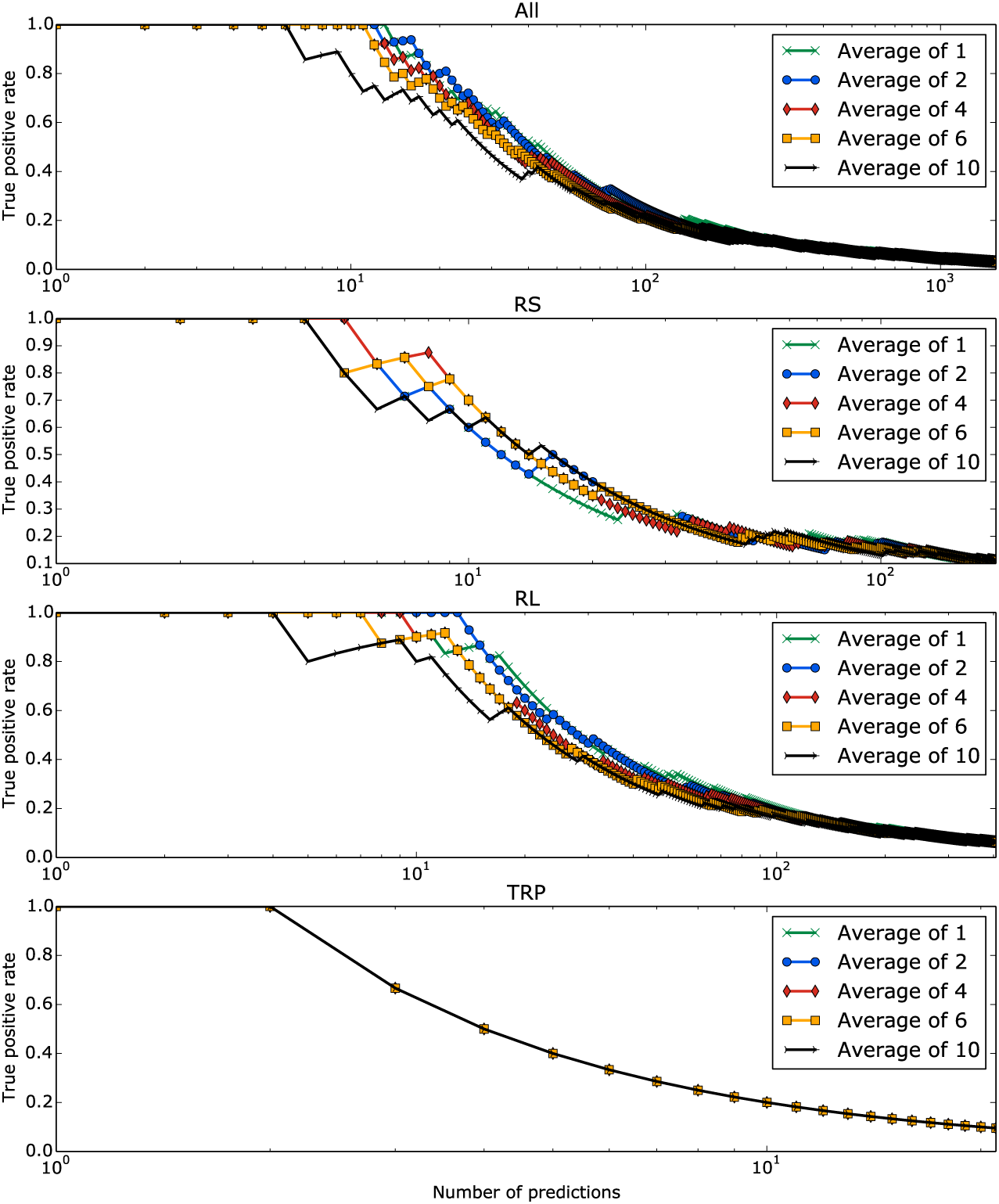
True positive rates at a given number of predictions; All: All possible protein pairs between RS, RL and Trp proteins are considered; RS: Protein pairs within the small ribosomal subunit; RL: Protein pairs within the large ribosomal subunit; Trp: Protein pairs of the Trp operon. Different lines indicate a different number of averaged inter-protein scores on the residue level to get a protein-protein interaction score

**Figure 16:**
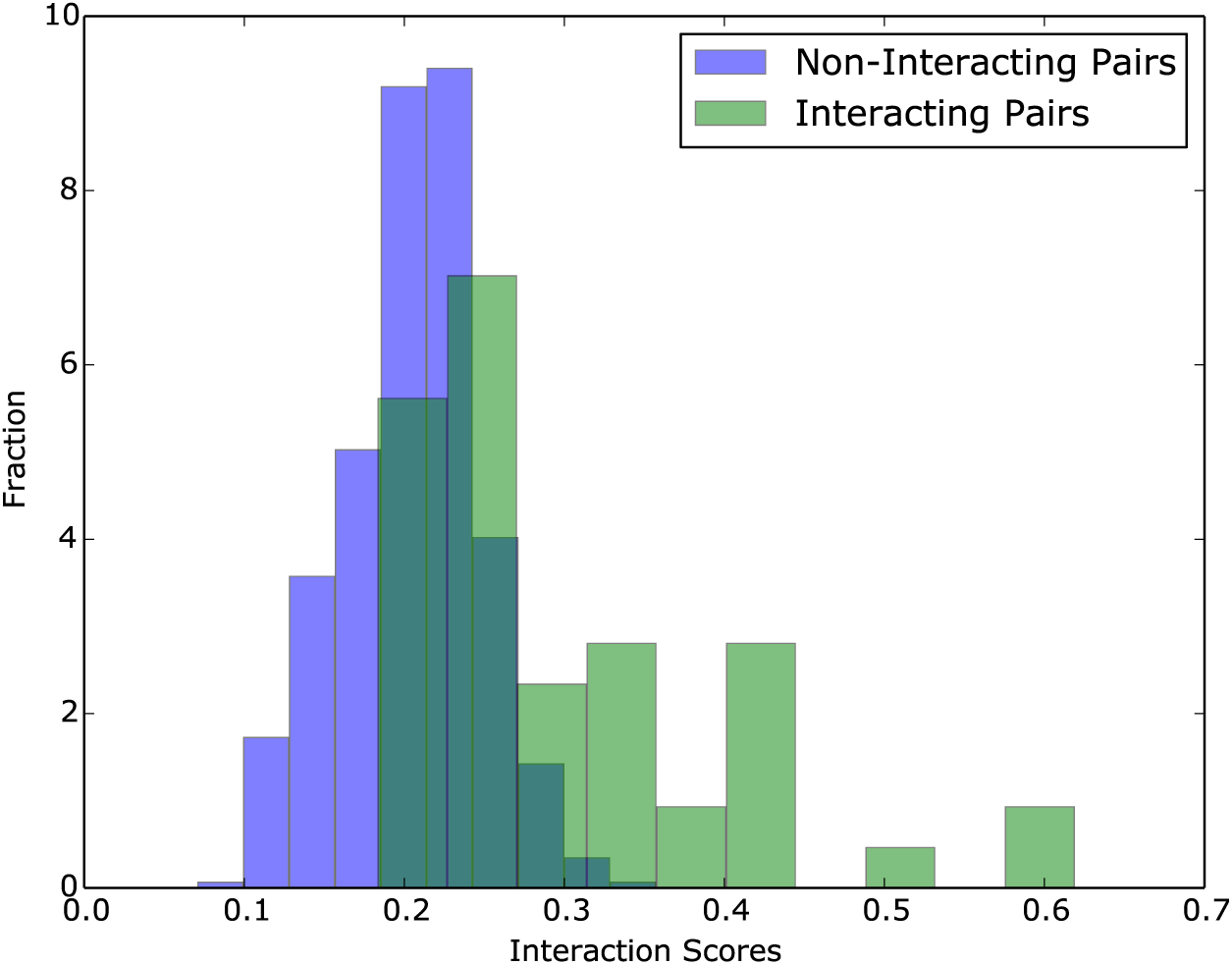
Histograms of interaction scores in the network comprising all possible protein pairs between RS, RL and Trp proteins. The protein-protein interaction scores were calculated averaging the 4 largest inter-protein residue interaction scores (as in the main paper). The histogram shows true positives and true negatives seperately. Both histograms are normalized.

